# A Grapevine MYC2-MYB24 Regulatory Module Activates Terpenoid Biosynthesis Upon Methyl Jasmonate Elicitation

**DOI:** 10.1101/2025.01.27.635049

**Authors:** Chen Zhang, Jone Echeverria, Chiara Foresti, Antonio Santiago, Hafsa El Idrissi Moubtassim, Alessandra Amato, Paolo Sonego, Massimo Pindo, Sara Zenoni, Marco Moretto, José Tomás Matus

## Abstract

Terpenes contribute to the characteristic flavor and aroma of grapes and their derived products while serving protective roles in plants against radiation, oxidative stress, and biotic challenges. The phytohormone methyl jasmonate (MeJA) mediates many of these processes and enhances terpene content in grape berries, but its underlying regulatory mechanism remains unclear. To address this, we treated *Vitis vinifera* cv. ‘Gamay Fréaux’ berry cell suspensions with MeJA (100 μM) and cyclodextrins (50 mM), generating transcriptomic data. Upregulated genes (URGs) were enriched in jasmonic acid biosynthesis and signaling pathways, and transcription factors (TF) that represent novel candidates modulating MeJA responses. Inspection of these TFs in terms of their co-expressed genes allowed us to focus on *MYC2*, the sole bHLH IIIe subgroup member in grapevine. Using DAP-seq, we identified MYC2-bound genes and integrated them with MeJA-URGs and *MYC2* co-expressed genes (CEGs) to produce a high-confidence target list. These targets were bound via a conserved G-box motif and included jasmonate-related genes (e.g., *LOX* and *JAZ*) and TFs such as MYB24, previously found to interact with MYC2 to activate terpenoid biosynthesis genes. Consistently, MeJA treatment induced 15 terpene synthase genes (*TPS*), eleven of which were bound by MYB24, MYC2, or both. Terpenoid compounds associated with these induced *TPSs* accumulated both intra- and extracellularly following treatment with MeJA and cyclodextrins, but not with abscisic acid (ABA). Our findings suggest that MYC2 regulates the jasmonate pathway and cooperates with MYB24 to mediate MeJA-induced terpene biosynthesis, shedding light on mechanisms that may extend to flower and fruit development, where this MYB-bHLH complex is also expressed.

## Introduction

Plants have developed different strategies to adapt and respond to the changes occurring in their environment. Some of these responses include chemical defense mechanisms, with the release of volatile organic compounds (VOCs) that are capable of diffusing in the air and trigger molecular and physiological responses far from their production site (i.e., in other organs or even in other plants). These molecules not only serve as a communication channel but also possess an effect on animals (e.g., repelling herbivore insects or attracting insect predators) and microorganisms (e.g., showing a negative effect on the growth of bacteria or fungi).

VOCs are represented by a vast group of molecules belonging to very different metabolic pathways including aldehydes, green leaf volatiles, terpenoids, among others. Within this group, terpenoids correspond to a large group of isoprenoid compounds synthesized both in the cytosol and chloroplasts, involved in a broad range of molecular processes such as growth regulation, herbivore defense, abiotic protection, animal attraction, and antimicrobial activity (Bosman and Lashbrooke, 2023). Among the most well studied biological functions, mono and sesquiterpenes play versatile roles serving as airborne signals that promote defense responses within plant communities (Shiojiri et al., 2015; Riedlmeier et al., 2017; Frank et al., 2021), attracting pollinators and natural enemies of herbivores (Arimura et al., 2004; Kappers et al., 2005; Schnee et al., 2006; Filella et al., 2013), and acting as antioxidants to protect plants from UV light and heat (Copolovici et al., 2005; Sasaki et al., 2007; Gil et al., 2012). In grapevine (*Vitis vinifera*), terpenes play a crucial role in shaping the aroma blend (also known with the French term ‘bouquet’) of table and wine grapes and their derived products, i.e., raisins and wines, respectively, thus contributing to their quality. Terpenes are essential for building the distinctive varietal characteristics of wines, so-highly valued in the market.

Terpene production is typically triggered during specific developmental stages, such as flowering, or in response to biotic stresses like herbivore attacks or pathogen infection. These processes are often mediated by the phytohormone jasmonic acid (Tholl, 2006; Yuan and Zhang, 2015; Rosenkranz et al., 2021). Its methyl ester, methyl jasmonate (MeJA), is widely used to elicit herbivore resistance in various plant species, leading to the production and release of terpenes (Baldwin, 1998; Li et al., 2002; Miller et al., 2005). MeJA has historically been used as an elicitor of terpene biosynthesis through exogenous applications on grapevine fruits (Gómez-Plaza et al., 2012; Ruiz-García et al., 2014; D’Onofrio et al., 2018; Li et al., 2020a; Marín-San Román et al., 2020; Wang et al., 2022; Zhang et al., 2024). Several studies have demonstrated that the induction of general isoprenoid and terpenoid biosynthetic genes is accompanied by an increase in berry terpene concentrations following MeJA treatments (Wang et al., 2022; Zhang et al., 2024; D’Onofrio et al., 2018). As an example, the transcriptomic study performed by Almagro *et al*., 2014 showed the term ’terpene biosynthesis’ to be significantly enriched among upregulated genes in cells treated with MeJA, cyclodextrins, and a combination of both. Despite this, the accumulation or release of terpenes from these elicited cells was never studied.

Recent studies in several plant species have shown that MeJA and cyclodextrin (CD) can elicit the production of specialized metabolites belonging to the phenylpropanoid pathway, such as resveratrol (Lijavetzky *et al*., 2008) and oxyresveratrol (Santiago et al., 2024). CDs refer to cyclic oligosaccharides composed of glucose residues linked by α-(1, 4)-glycosidic bonds, suggested to support the transport of MeJA and hydrophobic metabolites, facilitating their import and release, respectively, without compromising cell growth. They form complexes with specialized metabolites to reduce biosynthesis feedback inhibition, prevent degradation, and enhance stability (Pinho et al., 2014; Cardillo et al., 2021). Despite the combined use of MeJA and CD represents a promising strategy for enhancing metabolic yield, the effect of these elicitors on terpene production remains unexplored.

The biosynthesis of terpenes involves a complex pathway that begins with the formation of isoprene units. These units are then assembled into larger molecules through the action of enzymes known as terpene synthases (TPSs). A few studies have focused on the relative expression of *TPS* genes to deduce terpene production in response to MeJA (D’Onofrio et al., 2018; Wang et al., 2022; Zhang et al., 2024). Given that structural genes of specialized metabolic pathways are largely regulated by transcription factors, it stands to reason that TFs regulating *TPS* gene expression should also respond to MeJA. However, the mechanism under which MeJA induces *TPS* gene expression, with the subsequent terpene production, has not been studied yet.

Here, we applied a multi-omic approach to understand the regulation of terpene biosynthesis in grapevine using a cell suspension system derived from cv. ‘Gamay Freaux’ berries. We generated transcriptomic data from cells treated with methyl jasmonate (MeJA) and methyl-beta-cyclodextrin (CD). Global transcriptomic approaches were integrated with gene co-expression networks to narrow down the pool of potential transcription factors regulating terpene synthesis. Genome-wide inspection of transcription factor binding sites, coupled with experimental validation of TF activation and assessment of terpene production, allowed us to unveil a core regulatory module involving MYC2 and MYB24, which mediates the synergistic effects of MeJA and CD elicitation.

## Results

### Methyl jasmonate and methyl-beta-cyclodextrin stimulate JA signaling and terpene biosynthesis pathways

A *Vitis vinifera* cv. ‘Gamay Fréaux’ var. teinturier cell suspension system was employed to perform transcriptomic analysis, investigating the effects of methyl jasmonate (MeJA) and methyl-beta-cyclodextrin (CD) treatments on grapevine cells. The cells were treated with a combination of MeJA and CD (MeJA+CD) and harvested on day 4. To minimize technical variability within groups, 12 samples were sequenced and analyzed across both the treatment and control groups. Principal component analysis (PCA) revealed clear clustering of samples based on treatment conditions, with PC1 accounting for 84.42% of the variance, highlighting significant differences between elicited cells and control samples (Figure 1A). Differential expression analysis was performed to identify differentially expressed genes (DEGs) by comparing MeJA+CD-treated cells to control cells, using a significance threshold of *p* ≤ 0.05. DEGs were categorized into up-regulated genes (URGs, log2FC ≥ 1.5) and down-regulated genes (DRGs, log2FC ≤ -1.5) (Supplemental Table 1). Additionally, a publicly available transcriptomic dataset, generated under a similar experimental design where cells were treated with either CD, MeJA alone, or their combination, with samples collected at 24 hours, was re-analyzed (Almagro et al., 2014). For this dataset, DEGs were filtered using the same significance threshold (*p* ≤ 0.05), while URGs and DRGs were defined as having an absolute log2FC > 0.58, reflecting the microarray platform used to generate the data, which incorporates thousands of DNA probes (Supplemental Table 2).

**Figure 1.**
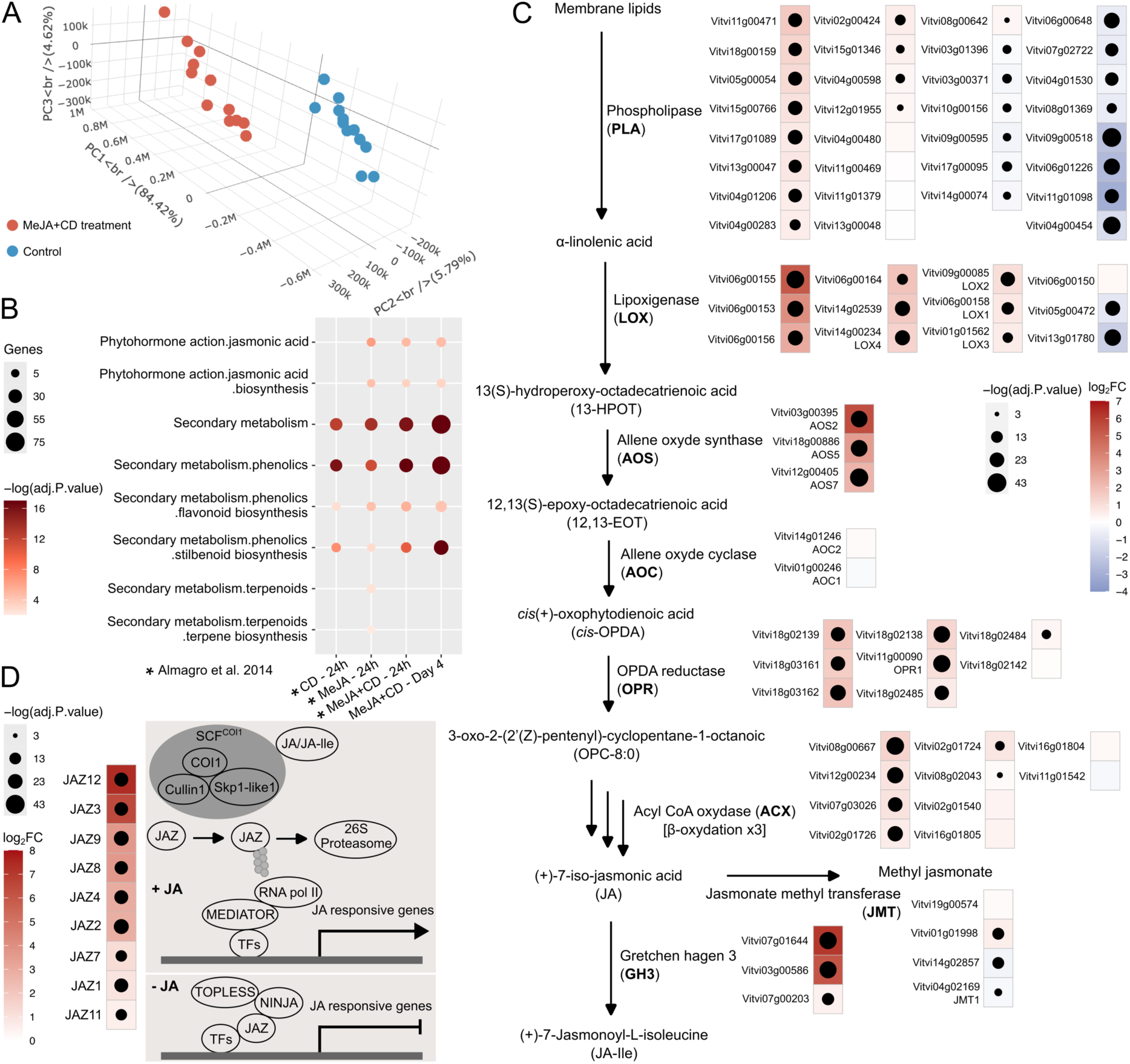
Activation of jasmonic acid (JA) and terpene biosynthesis pathways in cells elicited with MeJA and MeJA+CD treatments. (A) Principal component analysis (PCA) of transcriptomic data from MeJA+CD and control treatments. PCA shows clustering of samples based on treatment conditions, with PC1 accounting for 84.42% of the variance. (B) Enrichment analysis of jasmonic acid- and terpenoid-related terms in up-regulated genes (URGs) from MeJA and MeJA+CD treatments. Statistical significance is displayed on a –log(adj. P.value) scale, indicated by a color gradient from light red to deep red. Dot sizes represent the number of URGs intersecting with each ontology term. (C, D) Differential expression of genes associated with the JA biosynthesis and signaling pathways. Statistical significance is represented by a –log(adj. P.value) scale, with differential expression compared to controls shown as a color gradient from light blue to red (log2FC). Dot sizes indicate the level of significance.

To explore the positive cellular responses triggered by experimental treatments, up-regulated genes (URGs) were analyzed through gene set enrichment analysis (GSEA) using the MapMan ontology file. Enrichment terms related to flavonoid and stilbenoid biosynthesis were significantly enriched in the URGs of cells treated with MeJA, CD, and their combination (Figure 1B). Interestingly, the term ‘Secondary metabolism.terpenoids.terpene biosynthesis’ was significantly enriched only in the URGs of MeJA-treated cells (Figure 1B). To address the potential negative effects of CD, we examined the intersected URGs from MeJA+CD-treated cells associated with terpene biosynthesis. Seven URGs from MeJA+CD-treated samples collected on day 4 were found to intersect in the term ‘.terpene biosynthesis’, though with a non-significant adjusted p-value (Supplemental Table 3). In contrast, MeJA-treated cells contained five intersected URGs in this term, showing a significant adjusted p-value (Supplemental Table 3). This discrepancy could be explained by considering the query size, which represents the number of URGs annotated in the MapMan file and serves as the pool size for the analysis, directly influencing the p-value calculation (Supplemental Table 3; Supplemental Figure 1). Overall, these findings suggest that MeJA treatment, alone or in combination with CD, effectively induces terpene biosynthesis-related genes in grapevine cells.

We also observed that the term ‘Phytohormone action.jasmonic acid.biosynthesis’ was significantly enriched in the URGs when MeJA was used for elicitation (Figure 1B). To better understand how MeJA triggers the jasmonic acid (JA) biosynthesis pathway, the expression profiles of genes involved in this pathway were analyzed using transcriptomic data from MeJA+CD-elicited cells collected on day 4. The MeJA+CD treatment enhanced the expression of genes controlling each step of the JA biosynthesis pathway (Figure 1C). Previous studies have demonstrated that transcriptional changes in response to MeJA treatment in cell cultures primarily affect genes involved in JA signaling, such as JAZ and MYC transcription factors. Notably, JAZ repressors are among the earliest genes to be induced in response to JA treatments or wounding (Chini et al., 2007; Pauwels et al., 2009). In our analysis, transcript levels of nine JAZ genes were significantly upregulated (Figure 1D).

### MeJA and CD-induced MYC2 and MYB24 as regulators of JA and terpene biosynthesis Pathways

To decipher the regulatory mechanisms activated by MeJA+CD treatment at day 4, transcription factors (TFs) were identified among URGs using the Plant Transcription Factor Database (PlantTFDB, Jin *et al*., 2017) and manually confirmed through literature review. A total of 64 TFs were identified from 1,112 URGs, with 52 of them belonging to well-characterized families such as bHLH, NAC, bZIP, WRKY, AP2/ERF, MYB, and TIFY (Licausi et al., 2010; Zhang et al., 2012; Wang et al., 2013; Liu et al., 2014; Wang et al., 2014; Guo et al., 2016; Wong et al., 2016; Li et al., 2017; Wang et al., 2018) (Figure 2A; Supplemental Table 4). Using these family-characterized TFs, the TOP420 (1%) co-expressed genes (CEGs) were analyzed in a tissue-independent context via the VitViz platform (Supplemental Table 5). These CEGs were further examined using GSEA with the MapMan ontology file to explore their potential roles (Supplemental Table 6). Significant enrichment was observed in URGs for terms such as ‘Phytohormone action.jasmonic acid’, ‘Secondary metabolism.terpenoids’, and ‘Secondary metabolism.phenolics’ in MeJA- and MeJA+CD-treated cells. These terms were also identified among enriched terms of CEGs, allowing the construction of a network showing correlations between TFs and selected enriched terms of their CEGs (Figure 2B).

**Figure 2.**
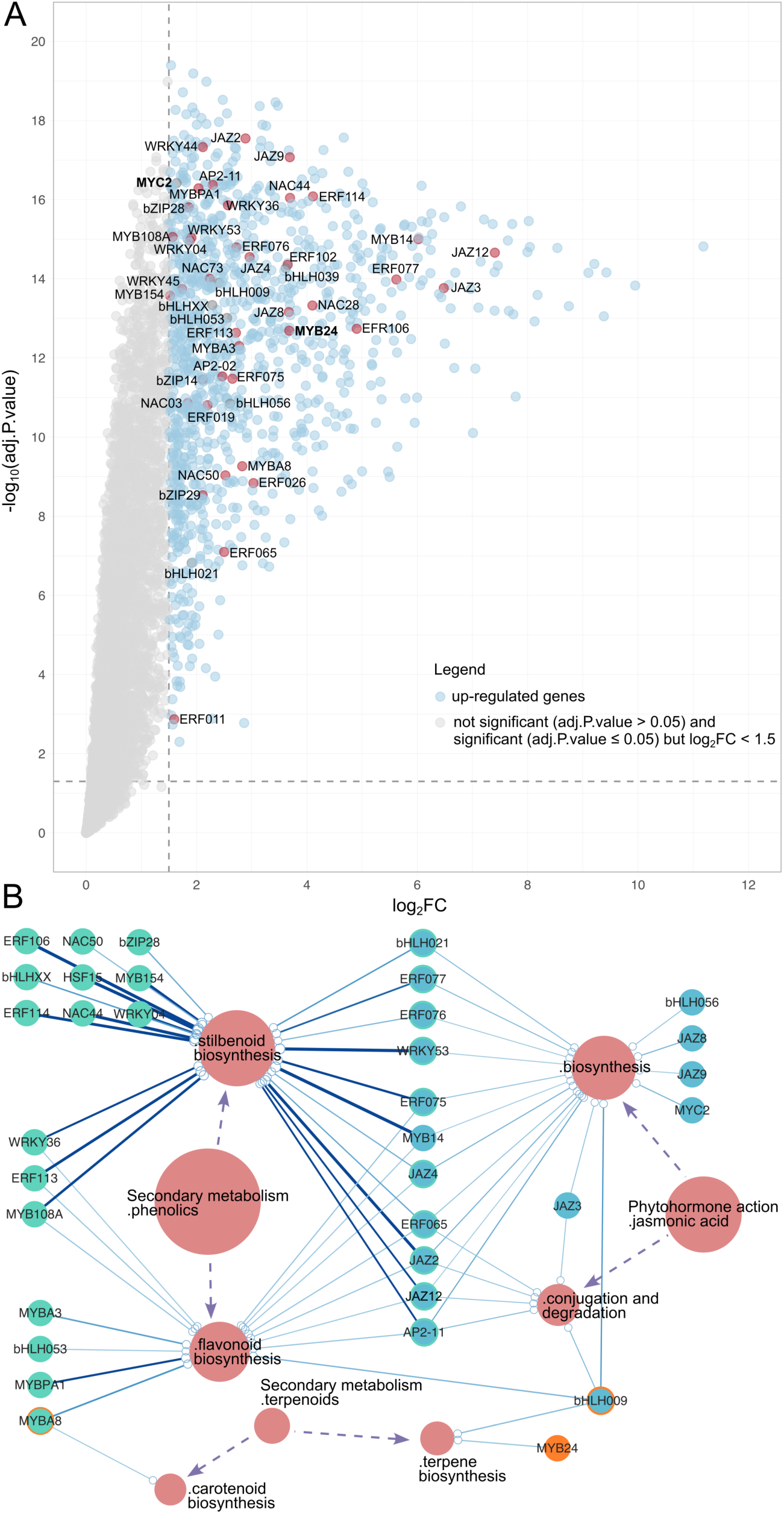
Regulatory roles of MeJA+CD-induced MYC2 and MYB24 in JA and terpene biosynthesis pathways. (A) Prediction of transcription factors (TFs) significantly induced in MeJA+CD-treated cells (logFC ≥ 1.5). Up-regulated genes are represented as blue dots, while transcription factors are marked by red dots. (B) A network visualization of MeJA+CD-induced transcription factors and a selection of enriched terms derived from their co-expressed genes (CEGs). Enriched terms are displayed as red nodes, with node sizes proportional to the number of connections with TFs. TFs are represented by nodes in orange, light green, or teal, indicating associations with ‘secondary metabolism.terpenoids’-related, ‘secondary metabolism.phenolics’-related, and ‘phytohormone action.jasmonic acid’-related terms, respectively. Nodes outlined with orange or light green circles indicate TFs associated with both ‘secondary metabolism.terpenoids’- and ‘secondary metabolism.phenolics’-related terms. The statistical significance of enriched terms, represented as edges, is displayed on a –log(adj. P.value) scale, with a color gradient ranging from light blue to deep blue. Edge widths correspond to the number of intersected co-expressed genes within the enriched terms. Purple dashed arrows indicate the hierarchical organization of MapMan ontology terms.

Notably, ‘Secondary metabolism.phenolics’ and ‘Phytohormone action.jasmonic acid’-related terms showed a larger number of intersected URGs and higher significance (adj.P.value) (Figure 1B). Approximately 63.5% of the family-characterized TFs were correlated with these terms (Figure 2B; Supplemental Table 6). Experimentally validated functions of TFs supported these findings. For example, MYB14 activates the promoter activity of *STS* genes, regulating stilbene biosynthesis in grapevine (Höll et al., 2013), while *MYBPA1* is involved in proanthocyanidin biosynthesis (Bogs et al., 2007). *MYC2*, the sole member of the IIIe bHLH subgroup in grapevine (Zhang et al., 2023), was associated with the term ‘Phytohormone action.jasmonic acid.biosynthesis’ (Figure 2B; Supplemental Table 6), and was upregulated in all MeJA-treated cells (i.e., MeJA and MeJA+CD; Supplemental Figure 2). In *Arabidopsis thaliana* and tomato, members of the IIIe bHLH subgroup have been extensively studied in JA signaling (Fernández-Calvo et al., 2011; Du et al., 2017; Chico et al., 2020). However, their roles in regulating the JA pathway in grapevine remain uncharacterized, making *MYC2* a strong candidate for investigating transcriptional regulation of the JA pathway in grapevine cell cultures elicited by MeJA.

Among TFs connected to ‘Secondary metabolism.terpenoids’, *MYBA8* was linked to ‘carotenoid biosynthesis’, while *MYB24* and *bHLH009* were associated with ‘terpene biosynthesis’ (Figure 2B; Supplemental Table 6). To confirm no additional MeJA+CD induced TFs were involved in regulating terpene biosynthesis, CEGs of 12 family-uncharacterized TFs were analyzed, but no significant enrichment was observed in the ‘terpene biosynthesis’ term (Supplemental Table 5 and 6). *VvibHLH009*, a member of the IIId bHLH subgroup in grapevine, is phylogenetically close to *Arabidopsis AtbHLH3*/*13*/*14*/*17*, which act as transcriptional repressors due to the absence of a conserved activation domain (Song *et al*., 2013; Zhang *et al*., 2023; Nakata *et al*., 2013; Fonseca *et al*., 2014). Sequence analysis of *VvibHLH009* revealed a similar non-conserved activation domain (Supplemental Figure 3). In contrast, MYB24, which has been shown to regulate terpene biosynthesis through activation of *TPS35* and *TPS09* in response to light in grapevine, under the participation of MYC2 (Zhang et al., 2023), displayed stronger induction in MeJA+CD-elicited cells (Figure 2A). Compared to *bHLH009*, *MYB24* had higher induction, while its partner *MYC2* was also significantly increased in MeJA+CD-elicited cells (Figure 2A). Its co-expression with MYC2 suggests a robust regulatory potential for activating terpene biosynthesis in response to MeJA+CD treatment.

### MYC2 regulates JA biosynthesis and signaling through hierarchical and conserved binding mechanisms

MYC2, a member of the grapevine bHLH transcription factor family, was localized in the nucleus of agroinfiltrated *Nicotiana benthamiana* leaves, confirming its expected subcellular location (Supplemental Figure 4). To gain a detailed understanding of MYC2’s regulatory landscape, DAP-seq was performed, enabling the analysis of its *in vitro* binding to genomic DNA from the Pinot Noir cultivar. This analysis identified 35,051 binding sites, which were mapped to their nearest gene features in the PN40024 reference genome, revealing a total of 15,783 target genes (Supplemental Table 7). Binding peaks were distributed across upstream, intragenic, and downstream regions of MYC2-bound genes, with a notable enrichment in upstream regions (Figure 3A). Interestingly, a slight decrease in MYC2 peak abundance was observed very close to transcription start sites (TSSs, Figure 3A).

**Figure 3.**
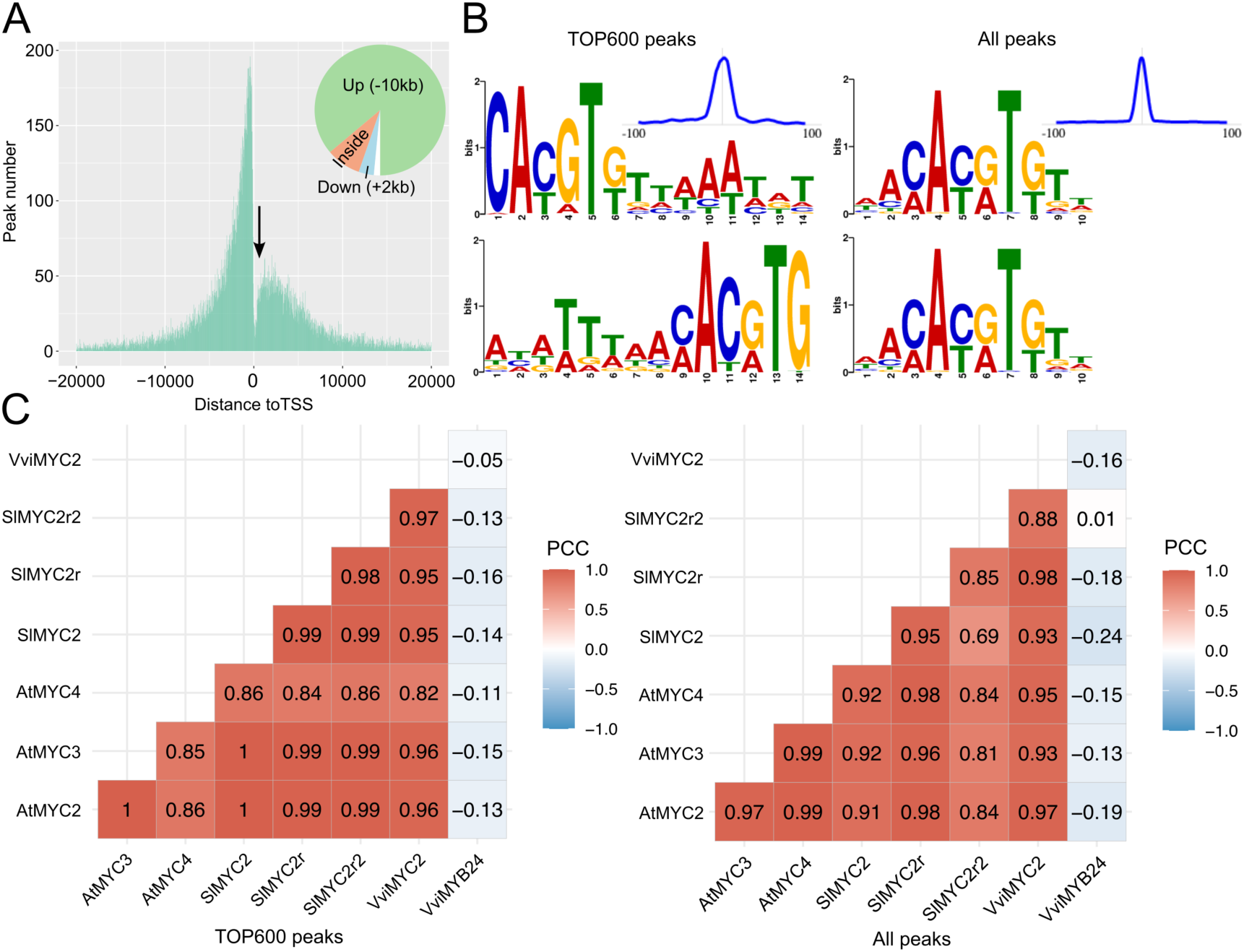
MYC2 consistently binds target genes through a conserved G-box motif. (A) Distribution of MYC2 binding peaks across the *Vitis vinifera* cv. ‘PN40024’ genome, relative to the transcription start sites (TSSs) of bound genes. Pie charts display the percentage of peaks located within specific genomic regions, ranging from ‘-10 kb/upstream’ to ‘+2 kb/downstream.’ An arrow highlights binding gaps near the TSSs. (B) Binding motif analysis from the 600 most significant peaks (left panel) and all detected peaks (right panel) in MYC2 DAP-seq data. Motifs are shown for forward (top) and reverse complement (bottom) strands. The blue curve in the upper-right corner represents the binding motif frequency profile across the analyzed peaks. (C) Cross-species comparison of consensus binding motifs identified in the top 600 peaks (left panel) and all peaks (right panel) among IIIe-bHLH members from *Arabidopsis thaliana*, *Solanum lycopersicum*, and *Vitis vinifera*. The consensus motif of the R2R3-MYB transcription factor VviMYB24, identified in the 600 most enriched peaks, served as a negative reference. Motif similarity, quantified by Pearson correlation coefficients (PCC), is represented on a color scale ranging from blue to red.

*De novo* motif analysis of the 600 most significant MYC2 binding peaks revealed a nearly identical core consensus sequence, ‘CACGTG’ (Figure 3B). This sequence corresponds to the G-box motif, a well-characterized DNA element recognized by bHLH proteins across plants, yeast, and animals (Ledent and Vervoort, 2001). Notably, flanking nucleotides such as thymidine (T) and adenine (A) residues were observed around the core G-box (Figure 3B), which were reported to modulate MYC2 binding activity (Dombrecht et al., 2007). To validate this consensus motif, we re-analyzed public DAP-seq data for IIIe-bHLH members from *Arabidopsis thaliana* and tomato, as generated by López-Vidriero *et al*., 2021. A similar consensus binding profile was identified among the top 600 peaks in these datasets, showing strong conservation of the G-box motif across *Arabidopsis thaliana*, tomato, and grapevine (Supplemental Figure 5). Among the seven IIIe-bHLH members analyzed, the G-box motif consistently emerged as the core binding sequence, with high similarity confirmed by Pearson correlation coefficients (PCC) (Figure 3C; Supplemental Figure 5).

The number of T-residues flanking the core G-box sequence is critical for its activation potential, as the removal of one or more T residues significantly reduces promoter activation (Kazan and Manners, 2013). Specifically, the presence of 3’-T nucleotides is essential for JA-dependent activation of *JAZ2* by MYC2, MYC3, and MYC4 (Figueroa and Browse, 2012). Given that approximately 50%, or in some cases more than half, of the genes were bound by one of the IIIe-bHLH members across the three species (Supplemental Table 8-13), we examined whether T-residues were consistently retained within the binding profiles of all detected peaks. Interestingly, the consensus binding motif for VviMYC2 showed a reduced number of T- and A-residues (Figure 3B), a pattern also observed for AtMYC3 (Supplemental Figure 5). In contrast, the consensus motifs of other IIIe-bHLH members retained extended T- and A-rich tails (Supplemental Figure 5). In addition to variations in flanking residues, subtle differences in the core binding sequences were observed among IIIe-bHLH members (Figure 3C; Supplemental Figure 5). While the G-box motif (CACGTG) remained the core binding sequence for VviMYC2 and *Arabidopsis thaliana* MYC2/3/4, as well as SlMYC2r, other members such as SlMYC2 and SlMYC2r2 displayed variants like ‘CACATG’ and ‘CATGTT’, corresponding to a G-box variant and a G-box-like sequence, respectively (Supplemental Figure 5). These findings suggest functional diversification within the IIIe-bHLH family, influenced by both core motif variants and flanking nucleotide composition.

To identify high-confidence targets (HCTs; Figure 4A; Supplemental Table 14) among MYC2-bound genes, we integrated DAP-seq data (binding sites spanning -5 kb upstream to 0.2 kb downstream of TSSs), *MYC2* co-expression data from the tissue-independent GCN (Supplemental Table 5) and MeJA+CD-induced genes (URGs at day 4; Supplemental Table 1). This approach yielded 698 HCTs (Supplemental Table 14), with a notable 14.8% predicted to encode transcription factors, as identified in PlantTFDB (Figure 4B). Among these, 6.3% were MeJA+CD-induced transcription factors (64 TFs; Supplemental Table 4), including members from key families such as MYB, NAC, WRKY, and TIFY. MapMan ontology analysis revealed that HCTs were enriched in pathways associated with ‘Secondary metabolism.phenolics.flavonoid biosynthesis,’ ‘Secondary metabolism.phenolics.stilbenoid biosynthesis,’ and ‘Phytohormone action.jasmonic acid.biosynthesis,’ consistent with patterns observed in URGs from MeJA+CD-treated cells (Figure 1B and 4C). Additionally, HCTs were significantly enriched in the term ‘RNA biosynthesis.Transcriptional regulation’ (Figure 4C), highlighting its regulatory roles. Among the HCTs, several genes involved in JA biosynthesis, including *LOX*, *AOS*, and *AOC*, were identified across the datasets (Figure 4D; Supplemental Table 14), along with *JAZ* (coding for members of the TIFY transcription factor family). These included *VviJAZ2*/*3*/*4*/*9* and *VviJAZ12* (Vitvi10g01879), a newly identified *JAZ* gene that was not originally classified in Zhang et al. (2012) (Figure 4A and E; Supplemental Table 14). Consistent with these findings, *Arabidopsis thaliana* AtMYC2 has been shown to positively regulate the expression of *AtJAZ* genes, directly binding to the promoters of *AtJAZ2* and *AtJAZ3* (Chini et al., 2007). Another noteworthy VviMYC2-bound target was *VvibHLH009*, a member of the IIId-bHLH subgroup (Figure 4A and E; Supplemental Table 14). In *Arabidopsis thaliana*, homologs of *bHLH009*, such as *AtbHLH017*, are negative regulators of JA responses and exhibit MYC2-dependent expression (Song et al., 2013; Fonseca et al., 2014). Altogether, these findings suggest that *MYC2* may orchestrate a complex transcriptional network of activation/repression processes to balance MeJA-induced responses.

**Figure 4.**
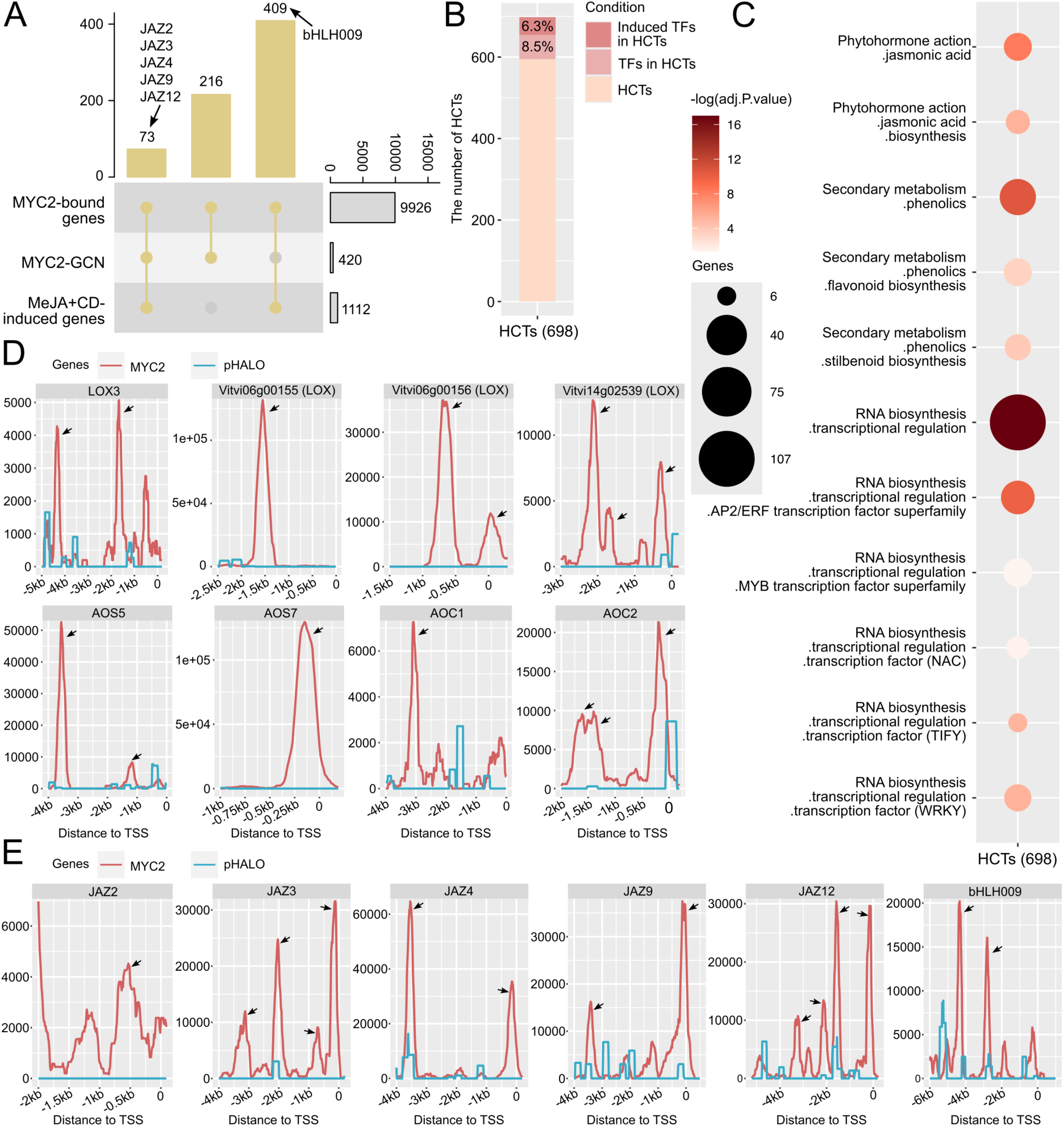
MYC2 mediates JA biosynthesis and signaling pathways through direct binding of their genes. (A) Definition of MYC2 high-confidence targets (HCTs), identified by overlapping three datasets: (i) MYC2-bound genes (filtered to include binding sites located from -5 kb upstream to 0.2 kb downstream of TSSs), (ii) MYC2’s co-expression network (GCN), generated using a tissue-independent aggregated network from the VitViz platform, and (iii) MeJA+CD-induced genes (log2FC ≥ 1.5). (B) Prediction of transcription factors (TFs) within the HCTs of MYC2. MeJA+CD-induced TFs are highlighted in red, non-induced TFs in the HCTs are marked in pink, and the remaining HCTs are indicated in light red. (C) Selected significantly enriched terms from MapMan ontology analysis of MYC2 HCTs. Statistical significance is represented on a –log(adj.P.value) scale, with a gradient from light red to deep red. The size of each dot reflects the number of URGs intersecting with each ontology term. (D) MYC2 binding signals at the promoters of JA biosynthetic genes (*LOX*, *AOS*, and *AOC*), compared to an empty vector (pIX-HALO) as a negative control. Arrows indicate detected binding peaks. (E) MYC2 binding signals at the promoters of JAZ genes and *bHLH009*, compared to an empty vector (pIX-HALO) as a negative control. Arrows indicate detected binding peaks.

### MYC2 and MYB24 complex promotes terpene accumulation in response to methyl jasmonate and methyl-beta-cyclodextrin

To investigate the transcriptional regulation of terpene synthesis in response to MeJA+CD elicitation, we analyzed terpene synthase genes (*TPS*) through differential expression analyses. Across two transcriptomic datasets—RNA-seq data from this study and public microarray data (Almagro et al., 2014)—a total of 31 *TPS* genes were found to be expressed (Supplemental Figure 6), 15 of which were significantly upregulated in elicited cells (Figure 5A). Building on prior observations that MYB24 and MYC2 exhibit strong regulatory potential in MeJA+CD-treated cells, we examined their binding capacities to these induced *TPSs*. MYB24 was previously shown to bind 22 *TPS* genes in a DAP-seq assay (Zhang et al., 2023). In the present study, MYC2 DAP-seq analysis revealed 113 distinct binding sites near 58 *TPS* genes (Supplemental Table 15). Notably, 11 *TPS* genes were both significantly induced under MeJA or MeJA+CD treatments and bound by either MYC2, MYB24, or both transcription factors (Figure 5A; Supplemental Figure 7).

**Figure 5.**
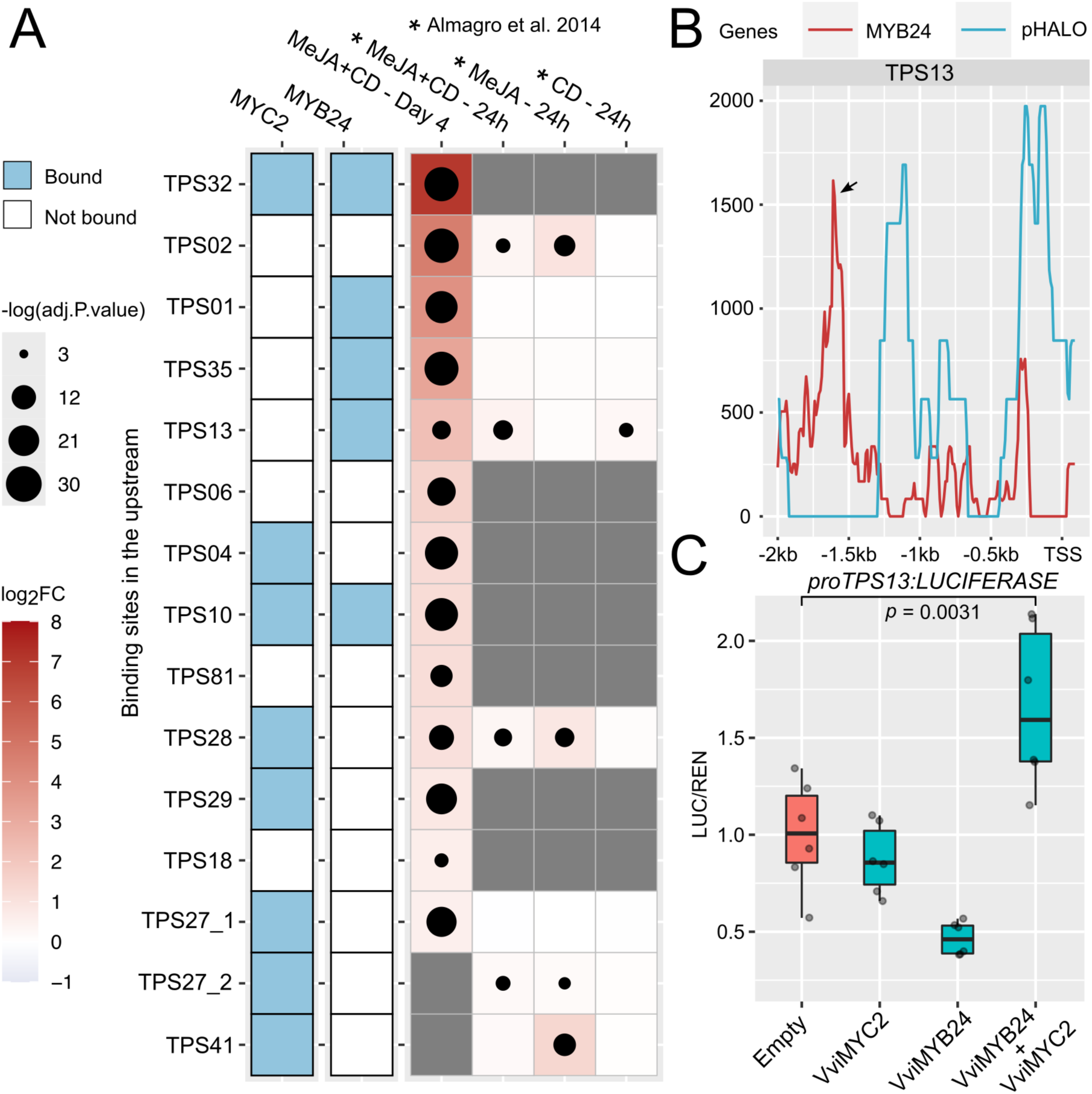
MYC2 and MYB24 complex activates terpene synthase genes (*TPSs*). (A) MeJA+CD-induced *TPSs* bound by MYB24 and/or MYC2. Left panel: *TPSs* bound by MYC2 and/or MYB24. Right panel: differential expression of *TPSs* in MeJA and MeJA+CD-elicited cells. Statistical significance is represented in a -log(adj. P.value) scale, with dot size reflecting significance. The color scale from light blue to red indicates differential expression compared with control samples (shown as log2FC). *TPSs* targeted by multiple DNA probes in the microarray are marked with underscores and numbers. (B) MYB24 binds to the promoter of *TPS13.* DAP-seq binding events were identified at -1.56 kb from the TSS (x-axis). Binding signals are compared with the empty vector (pIX-HALO) as a negative control. Arrows indicate detected binding peaks. (C) Transient expression of MYB24 and MYC2 activates *TPS13* promoter. Significance (p-values) was calculated based on one-way ANOVA followed by Tukey’s post hoc test. The boxplot shows the lower quartile, upper quartile, and median, with whiskers representing the minimum to maximum range.

Among the 11 MYC2- and/or MYB24-bound *TPS* genes, the binding distances for *TPS10*, *TPS13*, *TPS27*, and *TPS35* were within 5 kb upstream of their respective TSSs (Figure 5B; Supplemental Figure 7; Supplemental Table 15). Five of these TPS genes (*TPS01*, *TPS10*, *TPS13*, *TPS27*, and *TPS35*) have been functionally characterized through *in vitro* enzyme assays (Martin et al., 2010). TPS01, TPS13, and TPS27 produce the same terpene profile, consisting of 71% (*E*)-caryophyllene, 23% α-humulene, and 6% germacrene D, while TPS35 synthesizes β-ocimene, and TPS10 generates (*E*)-α-bergamotene. Considering their expression levels in MeJA+CD-elicited cells, proximity of binding sites, and confirmed terpene products, *TPS35* and *TPS13* were identified as prime candidates for further investigation into promoter activity influenced by MYB24 and MYC2. Activation of the *TPS35* promoter was previously demonstrated to require co-infiltration of MYB24 and MYC2 (Zhang et al., 2023). Similarly, a dual luciferase assay in *Nicotiana benthamiana* agroinfiltrated leaves showed enhanced *TPS13* promoter activity (spanning 1.56 kb upstream of the TSS) when MYB24 and MYC2 were co-infiltrated (Figure 5B and C).

As *TPS* gene induction was evident in elicited cells, we further quantified the corresponding terpene products synthesized by these induced *TPSs* (known roles of grape TPSs can be found in Supplemental Table 16). Terpenes are produced within cells and subsequently glycosylated (for storage) or released through the plasma membrane and cell wall, leading to their accumulation in the extracellular space. To account for this distribution, we quantified terpene levels in both elicited cells and in the growth media, representing intracellular and extracellular accumulation, respectively. A total of eight terpenes displayed significantly higher accumulation in the MeJA+CD treatment compared to the control, and these were detected in similar amounts in cells and their media (Figure 6). Among these terpenes, β-caryophyllene and α-copaene were quantified using available terpene standards, while the remaining terpenes were measured by peak area. MeJA and ABA alone did not elicit terpene production. Notably, six terpenes, including β-caryophyllene, α-copaene, (*Z*)-α-bergamotene, (*E*)-α-bergamotene, α-pinene, and 3-carene, also showed substantial synthesis in CD treatment compared with control samples (Figure 6). Despite this, extracellular and intracellular terpene accumulation was markedly more pronounced in the MeJA+CD treatment compared to single CD treatment, highlighting a synergistic effect of the combined elicitation (Figure 6).

**Figure 6.**
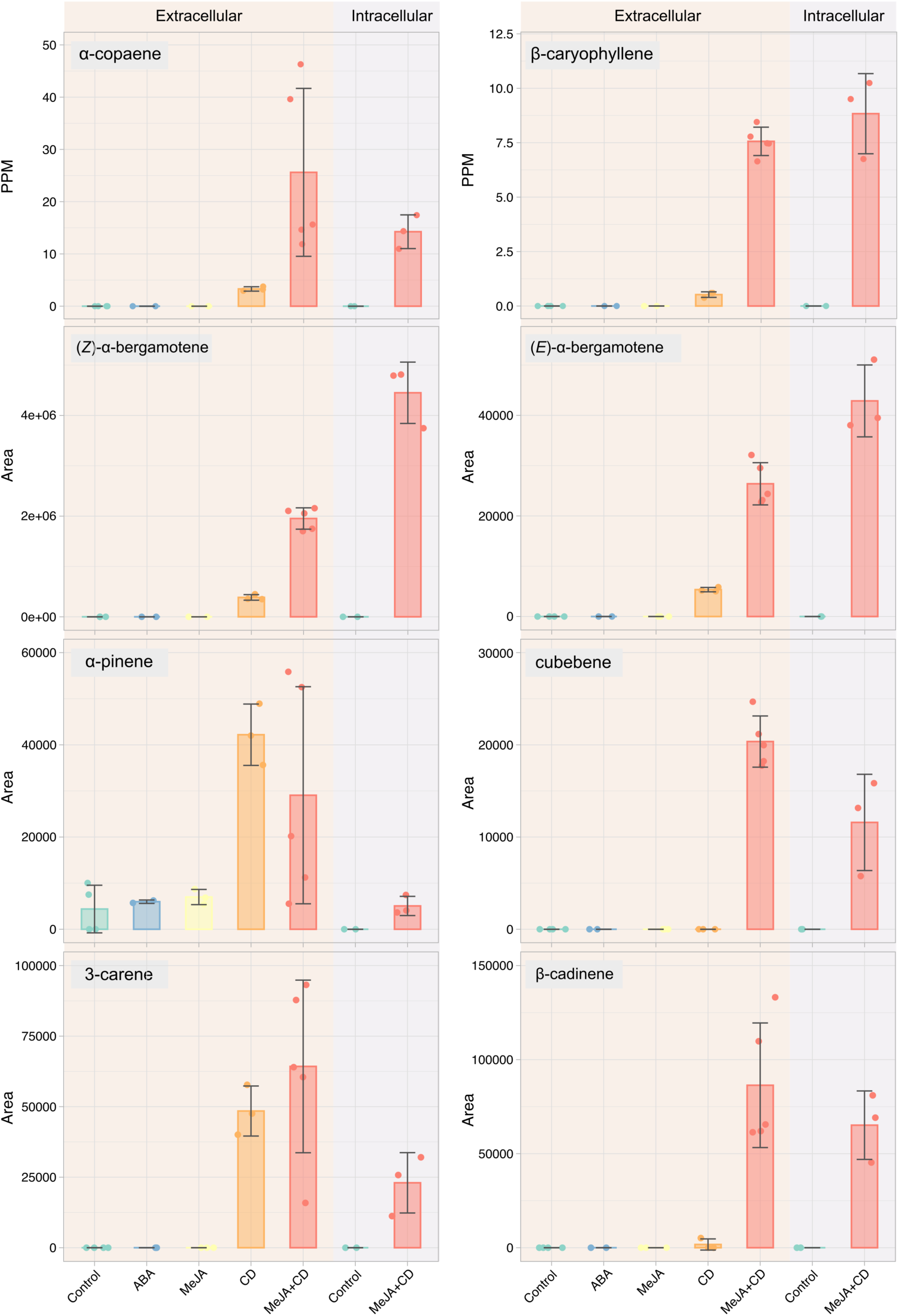
GC-MS targeted terpene quantification in response to MeJA and CD elicitation. Cells were treated with MeJA, CD, their combination, or ABA. Terpene levels were quantified in the treated cells and their respective media, representing intracellular and extracellular accumulation, respectively.

### Visualization of MeJA transcriptomic meta-analysis

To provide the research community with an exploratory tool for navigating MeJA-related responses in cell suspension experiments, we developed MeJA Atlas. This resource integrates gene expression data and DEGs from this study with reanalyzed data from the MeJA elicitation study by Almagro et al. (2014). Future datasets will be incorporated as they become available. MeJA Atlas, hosted on the PlantaeViz platform (https://plantaeviz.tomsbiolab.com/vitviz/meja_atlas/), enables interactive exploration using VitViz visualization tools. Users can input Vitis vinifera (Vcost) gene identifiers to generate heatmaps of differential gene expression. Each heatmap cell displays fold-change (indicated by color) and statistical significance (-log(p-value), represented by point size). This dual representation allows simultaneous assessment of expression magnitude and significance. The “Rep Heatmap” tab provides detailed visualization of expression patterns at the replicate level, enabling in-depth exploration of individual replicates.

## Discussion

Elicitation is an effective strategy for enhancing the production of specialized metabolites in plant organs and *in vitro* cell suspensions. This process is triggered by various elicitors, such as methyl jasmonate (MeJA) and cyclodextrin (CD) (Cardillo et al., 2021; Jeyasri et al., 2023). The major pathways of specialized metabolism known to be elicited by hormones in plant cell cultures correspond to phenylpropanoids and alkaloids (Cusido et al., 2014; Narayani and Srivastava, 2017). Here we show that structural enzymes of the sesqui and monoterpenoid pathway are also transcriptionally regulated in response to MeJA and CD elicitation. In addition, by integrating multi-omics approaches including transcriptomics, DAP-seq and GC-MS analysis, we identified *MYC2* as a central regulator of this MeJA-specific response. *MYC2*, the only member of the IIIe-bHLH subgroup within the grapevine bHLH transcription factor family, orchestrates jasmonic acid (JA) biosynthesis and signaling pathways in response to MeJA. Additionally, we show that MYC2 forms a protein complex with MYB24, which, in response to the combined effects of MeJA and CD elicitation, promotes terpene accumulation by activating terpene synthase genes (*TPSs*).

As demonstrated in this study and elsewhere, elicitation activates responses characteristic of plant defense. MeJA, the methyl ester of jasmonic acid, widely recognized as a key phytohormone mediating intra- and intercellular signaling in response to microbial pathogens and herbivores. Its high volatility enables MeJA to play a crucial role in organ-to-organ and plant-to-plant communication (Wang et al., 2021). The JA signaling pathway includes JAZ proteins being among the first genes to be induced (Larrieu and Vernoux, 2016). We demonstrated here that nine grape *JAZ* genes were significantly upregulated following MeJA and CD treatment. In *Arabidopsis*, these are largely known as MeJA-markers genes as they fine tune and act as negative feedback loops upon MeJA elicitation. The activation of the JA signaling pathway also necessitates the control of specific TFs, such as AtMYC2 allowing the subsequent responses of downstream genes (Pauwels and Goossens, 2011). Skp1/Cullin/F-box (SCF^COI1^) complex targets JA-related proteins (such as JAZs) for proteasomal degradation, but it requires a bioactive JA compound to recognize its substrates. While MeJA can be synthesized by jasmonate methyltransferase (JMT) within the JA biosynthetic pathway or applied externally to cells, it is not bioactive and cannot be recognized by the SCF^COI1^ complex. Among the various JA derivatives in plants, JA-Ile, which is synthesized by the GH3 family of enzymes, is the most bioactive and facilitates the SCF^COI1^ complex in targeting JAZs for degradation (Pauwels and Goossens, 2011; Wasternack and Strnad, 2016). Under internal or external stimuli, elevated JA-Ile promotes the degradation of SCF^COI1^-dependent JAZ inhibitors. Transgenic studies have demonstrated that MeJA must undergo cleavage and conversion to JA-Ile to become bioactive (Stitz et al., 2011). Our results showed that JA biosynthesis was positively regulated in MeJA and CD elicited cells. Notably, the expression of *GH3* genes was more strongly induced than *JMT*, suggesting an increased production of JA-Ile.

*MYC2*, the only member of IIIe-bHLH subgroup, was significantly upregulated by MeJA and MeJA+CD elicitation. Furthermore, high-confidence targets (HCTs) of MYC2 were significantly enriched in the gene ontology term ‘phytohormone action.jasmonic acid.biosynthesis’ and these overlapped HCTs included lipoxygenase (*LOX*), allene oxide synthase (*AOS*), and allene oxide cyclase (*AOC*) genes. This analysis also identified five MYC2-bound *JAZ* genes, including *JAZ2*, *JAZ3*, *JAZ4*, *JAZ9*, and *JAZ12*. In *Arabidopsis thaliana*, *AtMYC2* is a positive regulator of *AtJAZ* gene expression, and it has been proved that AtMYC2 directly binds to the promoter of *AtJAZ2* and *AtJAZ3* (Chini et al., 2007). Our results provide further evidence that MYC2 plays a crucial regulatory role in JA biosynthesis and signaling through direct binding of several JA-pathway genes. JAZ degradation, followed by MYC2-mediated transcriptional regulation and the resynthesis of JAZ repressors in response to MeJA stimulation, ensures that the activation and subsequent re-establishment of the JA signaling pathway occur in a tightly regulated manner. Additionally, *bHLH009*, a member of the IIId-bHLH subgroup, whose closest homolog (*AtbHLH017*) has been shown to negatively regulate JA responses in *Arabidopsis thaliana* (Fonseca et al., 2014), was also bound by VviMYC2 and induced in response to MeJA and CD treatment. These findings suggest that MYC2 acts as a key coordinator in the regulation of JA signaling pathway in response to MeJA elicitation, orchestrating a fine-tuned regulatory network.

Our results give support to the idea that cells produce secondary metabolites that play important roles in defense mechanism. In fact, the role of secondary metabolites, including flavonoids, terpenoids, and stilbenoids, in biotic and abiotic plant defense is well-documented across various plant species (Piasecka et al., 2015; Pérez-Pérez et al., 2024). It has been demonstrated that anthocyanins (i.e. flavonoids) and stilbenoids accumulate in cells elicited by MeJA and CD (Orduña et al., 2022). Consistently, we observed significant enrichment of gene ontology terms related to ‘Secondary metabolism.phenolics.flavonoid biosynthesis’ and ‘Secondary metabolism.phenolics.stilbenoid biosynthesis’ among the upregulated genes in MeJA and CD elicited cells. Additionally, 27 family-characterized TFs induced in MeJA and CD treated cells were associated with these enrichment terms. Particularly, these enriched terms and many of these family-characterized TFs appeared as analyzed results of HCTs of MYC2, suggesting that MYC2 may play a role in the transcriptional regulation of flavonoid and stilbenoid biosynthesis too. *In Arabidopsis thaliana*, AtMYC2 has been shown to form a regulatory module with *AtWRKY46* to positively regulate flavonoid accumulation as part of the E-2-hexenal-induced defense response (Hao et al., 2024). Further research is required to determine whether MYC2-mediated transcriptional regulation of flavonoid and stilbenoid biosynthesis in grapevine involves interactions with additional TFs.

Terpene quantification revealed significant accumulation under elicited conditions, with variations depending on the treatment. Eight terpenes accumulated at higher levels under the combined MeJA and CD treatment compared to MeJA or CD alone. This synergistic effect mirrors findings in other secondary metabolites, such as resveratrol and taxol, under similar elicited conditions (Lijavetzky et al., 2008; Cusido et al., 2014). Interestingly, while *TPSs* were strongly induced by MeJA treatment, only one was upregulated under CD treatment. Despite this, CD treatment alone yielded a higher quantity than MeJA, with six terpenes detected exclusively in CD-treated cells. This suggests CD may prevent terpene degradation and enhance their stability, similar to its proposed ‘protective’ role in stilbenoid stabilization (Silva et al., 2014). Complexation with CD and micellar systems has been shown to improve the solubility, stability, and bioactivity of stilbenes, which may also apply to terpenes. Although ABA has been reported to enhance the content of sesqui- and monoterpenes in grape leaves and berries (Murcia et al., 2017), we observed that ABA alone did not elicit terpene production in treated cells, consistent with our findings from the MeJA ‘alone’ treatment. The entry of ABA into cells is unlikely to be a limiting factor, as it is known to easily cross the cell membrane and be perceived by PYL receptors in the plasma membrane, making the combination with CD unnecessary. In fact, ABA in these cells promotes anthocyanin biosynthesis inside vacuoles demonstrating its effect (data not shown).

Among the 15 *TPSs* induced by the combined MeJA and CD treatment, five have been experimentally characterized through *in vitro* enzyme assays, specifically producing (*E*)-α-bergamotene, (*E*)-caryophyllene, α-humulene, germacrene D, and β-ocimene (Martin et al., 2010). Our results confirmed the accumulation of (*E*)-α-bergamotene and (*E*)-caryophyllene, synthesized by the induced TPS10 and TPS13, respectively. The remaining uncharacterized and induced TPSs may be responsible for the accumulation of the other detected terpenes. However, we cannot exclude the possibility that a single TPS may produce multiple terpenes in *planta*, despite showing specificity for certain compounds in both *in vitro* and *in vivo* assays (Martin and Bohlmann, 2004). Current literature suggests that TPS enzymes often exhibit ’promiscuous’ activity, lacking strict specificity for their products. Inconsistencies between the products generated in *in vivo* and *in vitro* conditions indicate that the enzymatic activity of a given TPS may vary depending on cultivars and environmental factors, such as substrate availability, climate conditions, and other external variables (Savoi et al., 2016). Additionally, allelic variations in *TPS* orthologs, resulting in amino acid substitutions across different cultivars, may further contribute to the diversity of terpene profiles observed (Smit et al., 2021).

Several studies have explored the transcriptional regulation of terpenoid biosynthesis in grapevine, identifying key transcription factors such as *WRKY40*, *ERF003*, *HYH*, and *GATA24* (Li et al., 2020b; Wang et al., 2024; Yang et al., 2024). Among the 64 TFs induced in MeJA and CD-elicited cells, *MYB24* was the only identified as a potential positive regulator of terpene biosynthesis during the elicitation process. In *Arabidopsis thaliana*, AtMYB24 and AtMYB21 are known to interact with bHLH transcription factors (AtMYC2/AtMYC5) to fulfill different roles in flower maturation (Qi et al., 2015). In grapevine, the cooperation between MYB24 and MYC2 was also validated for the activation of *TPS35* and *TPS09* promoters (Zhang et al., 2023). Among the induced *TPSs* bound by MYB24 and/or MYC2, we confirmed that activation of the *TPS13* promoter specifically required the presence of both MYB24 and MYC2, leading to the accumulation of the corresponding terpene product: β-caryophyllene. *MYC2* was indeed detected among the TFs induced in MeJA and CD-elicited cells, supporting the hypothesis that a regulatory scenario driven by the MYB24-MYC2 complex occurs during the elicitation process.

Our findings demonstrate that MYC2, as a central regulator, orchestrates JA biosynthesis and signaling pathways in response to MeJA. Furthermore, MYC2 collaborates with MYB24 to enhance terpene accumulation under the synergistic effect of combined MeJA and CD elicitation. The expression and function of MYC2 and MYB24 in the suspension cell system can be readily extrapolated to the physiological context of flower and berry development. These genes are strongly induced in these organs at distinct developmental stages and show a high correlation with *TPS* and jasmonate-related genes, as revealed by flower and fruit aggregated gene co-expression networks (Orduña *et al*., 2023; Supplemental Figure 8). Jasmonates are well known to promote flower maturation in various plant species, including *Arabidopsis thaliana*, where *MYC2* and *MYB24* homologs, as well as *TPS* genes, are highly expressed and respond strongly to exogenous MeJA treatments (Reeves et al., 2012; Qi et al., 2015). Similarly, in grapevine, *MYB24* and *TPSs* genes are expressed during late flowering, a stage associated with the accumulation of sesquiterpenes and monoterpenes (Zhang et al., 2023). This suggests that terpene biosynthesis in grape flowers is regulated by jasmonates. The elevated expression of *MYC2*, along with jasmonate biosynthesis and signaling genes, at the onset of berry ripening aligns with the hormone’s established role in this process in grapevine (Kondo and Fukuda, 2001; Jia et al., 2016). Additionally, during late berry ripening, the expression of MYC2, MYB24, TPS, and jasmonate-related genes indicates that jasmonates may regulate key molecular events at this stage, including the accumulation of terpenes, which are also known to peak late in ripening (Zhang et al., 2023). However, the roles of MYC2 and MYB24 in flower maturation and late ripening in grapevine remain speculative and require further validation within the plant context.

## Methods

### Plant material

*Vitis vinifera* cv. ‘Gamay Fréaux’ var. teinturier calli was used for the elicitation experiment. Solid calli was kept at 22°-24°C and dark condition, and was maintained in the media containing gamborg basal media (Duchefa), 2% sucrose, 0,205 g/L Bacto™ Casamino Acids (Gibco), 0,01 g/L Myo-inositol (Duchefa) and Vitis vitamin mix, and 7,5 g/L of agar. Subculture of solid calli was carried out every 3 weeks. Solid calli was transferred to 500 ml erlenmeyers with liquid media, generating the liquid culture and developing the cell suspension system. Liquid culture was placed in an orbital shaker with 110 rpm, and was grown at 20-24°C and dark conditions. Subculture of liquid culture was carried out every two weeks. *Nicotiana benthamiana* plants were grown in 25 °C (light, 18h) and 20 °C (darkness, 6h) condition.

### Methyl jasmonate and cyclodextrins treatment in cell suspension system

Seven-day old cells were used in the methyl jasmonate and cyclodextrins treatment. Liquid cells were filtered using sterile glass filters with 90-150 μm pores. Eight grams of cells were transferred to 32 grams of a new liquid media containing 100 μM methyl jasmonate (CAS No.1211-29-6) and 50 mM methyl-beta-cyclodextrin (MβCD, CAS No. 128446-36-6). Considering that methyl jasmonate was dissolved in methanol (100%), the same amount of methanol (100%) was used in the control samples. Liquid cells were filtered and collected at days 4 and 5 after treatment.

### Total RNA extraction

Total RNA was extracted from cell samples collected at day 4, which was following the instruction of Spectrum™ Plant Total RNA Kit (Sigma-Aldrich, USA). RNA samples were treated with DNase for removing gDNA contamination, following the instruction of DNA-freeTM Kit DNase Treatment and Removal Reagents (InvitrogenTM). Total RNA extracted from 4-days cell samples were used for generating transcriptomics.

### Transcriptomic analysis

Single-end reads of 100 bp were obtained for each of the 24 samples, comprising 12 control and 12 MeJA-treated conditions, using the Illumina NovaSeq 6000 sequencing platform. Approximately 30 million strand-specific sequences were generated per sample. The quality of the raw data was assessed using FastQC (version 0.11.9) (Leggett et al., 2013). The reads were preprocessed to remove adapters and perform quality trimming using fastp (version 0.23.2) (Chen et al., 2018). The preprocessed reads were aligned to the 12X.2 reference genome assembly of *Vitis vinifera* PN40024 using the STAR aligner (version 2.7.8a) (Dobin et al., 2013). Raw read counts were extracted with the featureCounts read summarization program (version 1.6.4) (Liao et al., 2014) based on the VCOST.v3 annotation. Genes with sufficiently large counts were retained using the edgeR (version 3.36.0) function filterByExpr following the filtering strategy described in Chen et al. 2016. Differential expression analysis was conducted using the limma empirical Bayes procedure (version 3.40.2) (Ritchie et al., 2015) with the limma-trend method (Law et al., 2014).

### Network generation

Transcription factors up-regulated in MeJA and CD-treated cells were predicted by PlantTFDB (Jin et al., 2017). TOP420 (1%) co-expressed gene list of each transcription factor was retrieved from aggregated gene co-expression networks (tissue-independent) (Orduña et al., 2023). Co-expressed gene of each transcription factor was analyzed by MapMan ontology file and the enriched terms that the number of intersected genes were smaller or equal to 2 were discarded. The selected terms significantly enriched in co-expressed genes of each transcription factor were visualized using Cytoscape v3.10.2.

### DNA affinity purification sequencing

DNA affinity purification sequencing (DAP-seq) of MYC2 was performed as described previously (Orduña et al., 2022; Fontanet-Manzaneque et al., 2024). *De novo* motif discovery was performed by retrieving 200 bp sequences centered at GEM-identified binding events for the 600 most enriched peaks and all peaks running STREME in MEME suit 5.5.3 with most default parameters. In total, if the number of peaks in DAP-seq analysis is more than 20000, the first 20000 peaks based on the q value are considered as all peaks to detect the consensus binding motif. The motif similarity was analyzed in R with ‘PWMSimilarity’ function of TFBSTools package.

### Volatile compound quantification

Cells and media collected at day 5 were used for volatile quantification. Frozen samples (400 mg of filtered cells and 400 μL of incubation media) were heated at 30°C for 10 minutes and 1.1 mL of a 5M CaCl2 solution and 200 µL of EDTA 500 mM, pH 7.5 were added. After gentle mixing, samples were subjected to headspace solid phase microextraction (HS-SPME).

A 65 uM PDMS/DVB (Supelco, Bellefonte, PA, USA) fiber was used for the analysis. Pre-incubation and extraction times were performed at 50 °C for 10 and 20 min respectively. Desorption was performed for 1 min at 250 °C in splitless mode. VOCs trapped on the fiber were analyzed by GC-MS using an autosampler COMBI PAL CTC Analytics (Zwingen, Switzerland), a 6890N GC Agilent Technologies (Santa Clara, CA, USA) and a 5975B Inert XL MSD Agilent, equipped with an Agilent J&W Scientific DB-5 fused silica capillary column (5%-phenyl-95%-dimethylpolysiloxane as stationary phase, 60 m length, 0.25 mm i.d., and 1 mm thickness film). Oven temperature conditions were 40 °C for 2 min, 5 °C min -1 ramp until 250°C and then held isothermally at 250 °C for 5 min. Helium was used as carrier gas at 1.4 ml min-1 constant flow. Mass/z detection was obtained by an Agilent mass spectrometer operating in the EI mode (ionization energy, 70 eV; source temperature 230 °C). Data acquisition was performed in scanning mode (mass range m/z 35–220). Chromatograms and spectra were recorded and processed using Agilent Masshunter software. Compound identification was based on both the comparison between the MS for each putative compound with those of the NIST 2017 Mass Spectral library and the match to their custom library generated using commercially available compounds.

### Dual-luciferase assay

The upstream region (promoter/5’ UTRs, 1783bp) of TPS13 was cloned into the entry vector pDONR. TPS13 promoter and the coding sequences of MYB24 and MYC2 were inserted into the reporter (pPGWL7) and effector vectors (pB2GW7) respectively by using LR clonase. Recombinant plasmids and the reference vector expressing the Renilla luciferase gene were transformed in *Agrobacterium tumefaciens* strain C58 by heat shock. Dual-luciferase assays were performed in *Agrobacterium* infiltrated leaves of *N. benthamiana* as described previously (Zhang et al. 2023).

### Web application

A dedicated web application (https://plantaeviz.tomsbiolab.com/vitviz/meja_atlas/) was developed as a Vitis Visualization (VitViz) tool within the PlantaeViz platform. This interactive tool allows users to explore normalized gene expression data and DEA results from this study, along with reanalyzed data from Almagro et al. (2014). Hosted on a Shiny server, the application offers a robust and scalable framework for deploying R-based web tools.

## Data availability

RNA-seq data generated in this study has been deposited at the NCBI Sequence Read Archive (SRA) under accession number PRJNA1213010. The roles for MYC2 and MYB24 have been deposited in the Gene Reference Catalogue found at the Grape Genomics Encyclopedia portal (http://grapedia.org/).

## Author contributions

J.T.M., C.Z, and J.E. designed the research. C.Z., J.E., C.F., H.E.M., A.A., and S.Z. conducted experiments. C.Z., A.S., P.S., M.P., and M.M. analyzed data. C.Z. and J.T.M. wrote the manuscript. All authors have read and approved the final version of the manuscript.

## Acknowledgements

We thank Ana Espinosa from the Metabolomics Service at the Institute for Plant Molecular and Cell Biology (IBMCP, Valencia) for her assistance and discussions in volatile analysis.

