## Supplemental Figures for "A Grapevine MYC2-MYB24 Regulatory Module Activates Terpenoid Biosynthesis Upon Methyl Jasmonate Elicitation"

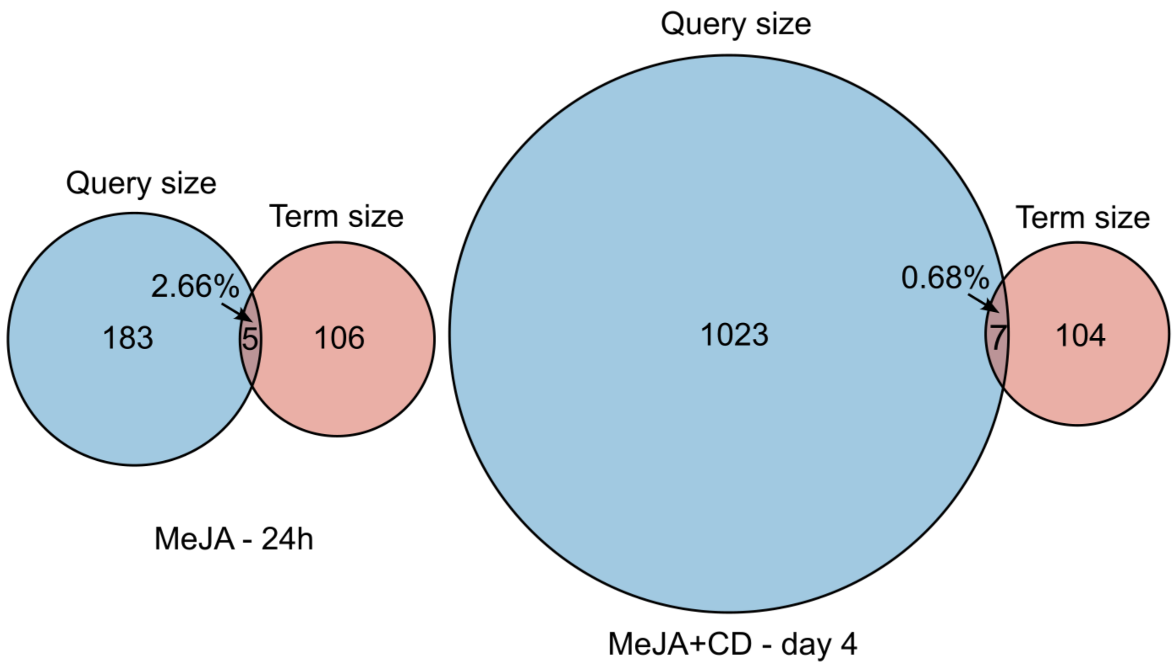


Supplemental Figure 1. Inspection of the intersection between up-regulated genes (URGs) detected in MeJA (at 24h; Almagro et al., 2014) and MeJA+CD (at 4 days; this study) treatments and genes belonging to the MapMan term "Secondary metabolism.terpenoids.terpene biosynthesis". Detailed analysis is shown in Supplemental Table 3.

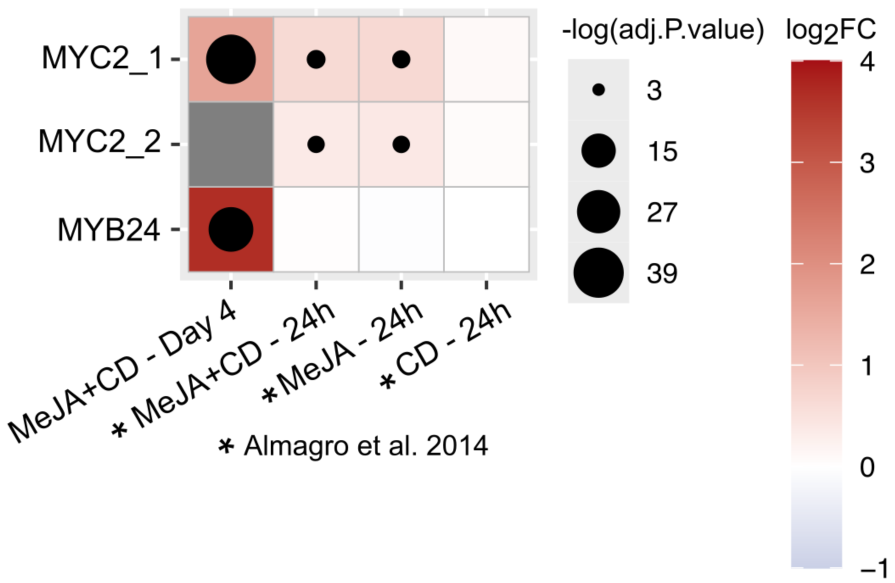


Supplemental Figure 2. The relative expression of *MYC2* and *MYB24* in MeJA- and MeJA+CD-elicited cells is depicted. Statistical significance is represented on a -log(adj. P-value) scale, indicated by dot size, while a color scale from white to red reflects differential expression compared to the control (log2FC). DNA probes on the microarray slide are denoted with an underscore.

```
AtbHLH03      ------------------MGQKFWENQE--DRAMVESTIGSEACDFFISTASASNTA-------LSKLVSP-----PSDSNLQQGLRHVV-----EGSDWDYALFWLASNVNSSD-----   78
AtbHLH013     ----------------MNIGRLVWNE-D--DKAIVASLLGKRALDYLLSNSVSNAN----------LLMTL-----GSDENLQNKLSDLVERPNASNFSWNYAIFWQISRSK-AG-----   80
AtbHLH017     ----------------MNMSDLGWDD-E--DKSVVSAVLGHLASDFLRANSNSNQN----------LFLVM-----GTDDTLNKKLSSLVDWPNSENFSWNYAIFWQQTMSR-SG-----   80
AtbHLH014     ---------------------------------MYN---------LTFSPSLSSSLLSFTQQTPAAIVSSSP-----PDLVLQQKLRFVVET---SPDRWAYVIFWQKMFDDQSD-----   65
VvibHLH009    ---------------------------------------------------------------------------------------------------------------------   0
AtMYC3        -------------MNGTTSSINFLTSDDDASAAAMEAFIGTNHHSSLFPPP-----------------PQQPPQPQFNEDTLQQRLQALIES---AGENWTYAIFWQISHDFDSST--GD   85
AtMYC4        MSPTNVQVTDYHLNQSKTDTTNLWSTDD--DASVMEAFIGGGSDHSSLFPP-----------------LPPPPLPQVNEDNLQQRLQALIEG---ANENWTYAVFWQSSHGFAGEDNNNN   98
AtMYC2        -------MTDYRL----QPTMNLWTTDD--NASMMEAFMSSSDISTLWPPASTTTTTAT-TETTPTPAMEIPAQAGFNQETLQQRLQALIEG---THEGWTYAIFWQPSYDFSGA-----   98
VviMYC2       -------MTEYRV-----PTMNLWT-DD--NASMMEAFISSDLSSFSWGPSSAASTSTPAPDPSRNLAQSQPSMAVFNQETLQQRLQALIEG---ARESWTYAIFWQSSVDFSGA-----   97

AtbHLH03      -GCVLIWGDGHCRVKKGAS-------------GEDYSQQDEIKRRVLRKLHLSFVGSDEDHRLVKSGALTDLDMFYLASLYFSFRCDTNKYGPAGTYVSGKPLWAADLPSCLSYYRVRSF   184
AtbHLH013     -DLVLCWGDGYCREPKEGEKSEIVRI---LSMGREEETHQTMRKRVLQKLHDLFGGSEEENCALGLDRVTDTEMFLLSSMYFSFPRG--EGGPGKCFASAKPVWLSDVVNSGSDYCVRSF   194
AtbHLH017     -QQVLGWGDGCCREPNEEEESKVVRSYNFNNMGAEEETWQDMRKRVLQKLHRLFGGSDEDNYALSLEKVTATEIFFLASMYFFFNHG--EGGPGRCYSSGKHVWLSDAVNSESDYCFRSF   197
AtbHLH014     -RSYLVWVDGHFCGNKNNNSQENYTT-------------------NSIECELMMDG-------------GDDLELFYA----ASFYGE--DRSPRKEVSDESLVWLTGPDELRFSNYERAK   147
VvibHLH009    ---------------------------------------------------------------------MDGDVATWEWYYMVSVTKSFVVG--DGVLGRVFSSGAFVWLTDR-ELQCYDCERVT   53
AtMYC3        NTVILGWGDGYYKGEEDKEKKK--NN--------TNTAEQEHRKRVIRELNSLISGGIGVSDESNDEEVTDTEWFFLVSMTQSFVNG--VGLPGESFLNSRVIWLSGSGALTGSGCERAG   193
AtMYC4        NTVLLGWGDGYYKGEEEKSRKKKSNP--------ASAAEQEHRKRVIRELNSLISGGVGGGDEAGDEEVTDTEWFFLVSMTQSFVKG--TGLPGQAFSNSDTIWLSGSNALAGSSCERAR   208
AtMYC2        --SVLGWGDGYYKGEEDKANPRRRSSS----PPFSTPADQEYRKKVLRELNSLISGGVAPSDDAVDEEVTDTEWFFLVSMTQSFACG--AGLAGKAFATGNAVWVSGSDQLSGSGCERAK   210
VviMYC2       --SLLGWGDGYYKGEEDKGKRKMTP---------SSVSEQEHRKKVLRELNSLISGTASSSDDAVDEEVTDTEWFFLVSMTQSFVNG--AGLPGQALFNSSPVWVVGTERLMSSPCERAR   204
                                                                                        : :        *                 :*  .           *

AtbHLH03      LARSAGFQTVLSVPVNSGVVELGSLRHIPEDKSVIEMVKSVFGGSDFV------------------QAK-------------------------------EAPKIFGRQLSL--------   247
AtbHLH013     LAKSAGIQTVVLVPTDLGVVELGSTSCLPESEDSILSIRSLFTSSLPPVRA----VALPVTV-AEKIDD-------------------------------NRTKIFGKDLHN--------   270
AtbHLH017     MAKSAGIRTIVMVPTDAGVLELGSVWSLPENIGLVKSVQALFMRRVTQPVM--------VTS-NTNMTG-------------------------------GIHKLFGQDLSG--------   269
AtbHLH014     EAGFHGVHTLVSIPINNGIIELGSSESIIQNRNFINRVKSIFGSGKTTKHTN-QTGSYP---------------------------------------------------------KPA---   208
VvibHLH009    EARMNGIRTLLCVSTSCGVLELGSLDMIKEDWGLVLLAKSLFGSKPS---------------------------------TQVSQ----------------IQIPDRNLSIFDIGAASG   123
AtMYC3        QGQIYGLKTMVCIATQNGVVELGSSEVISQSSDLMHKVNNLFNFNNGGGNNGVEASSWGFN-LNPDQGENDP-ALWISEPTNTGI--ESPARVNNGNNSNSNSKSDSHQISKLEKND---   306
AtMYC4        QGQIYGLQTMVCVATENGVVELGSSEIIHQSSDLVDKVDTFFNFNNGGGEF----GSWAFN-LNPDQGENDP-GLWISEPNGVDSGLVAAPVMNNGGN-DSTSNSDSQPISKLCNGS---   318
AtMYC2        QGGVFGMHTIACIPSANGVVEVGSTEPIRQSSDLINKVRILFNFDGGAGDLS--GLNWNL---DPDQGENDP-SMWINDPIGTPGSNEPGNGA-----PSSSSQLFSKSI-QFENGSSST   318
VviMYC2       QAQVFGLQTMVCIPSANGVVELGSTELIYQSSDLMNKVRVLFNFNNLE------VGSWPIGAAAPDQGESDPSSLWISDPTSNVEIKDSVNATATGASNPIGNQQNSKSI-QFENPSSSS   317
                .   *.:*:  :    *::*:**   : :. . :     .*

AtbHLH03      GGAKPRSMSIN-------------------------FSP-------------------KT----------------EDDTGFSL-------ESYEV---------------------   276
AtbHLH013     SGFLQHHQHHQQQQQQPPQQQQHRQFREKLTVRKMDDRAPKRLDAYPNNGN-RFMFSNPGT----------------NNNTLLSPT--WVQPENYTRPINVKEVPSTDEFKFLPLQQSSQ   371
AtbHLH017     AHAYPKK------------LEVRRNLDER--------FTPQSWEGYNNNKGPTFGYT-PQR----------------DDVKVLENVNMVVDNNNYKT----------------------   329
AtbHLH014     VS-----------DH--------------------------------------------------------------SKSGNQQFGSER-KRRR-KL-------------------   230
VvibHLH009    VQRE---------SH------------------------------------------------------------EGKQQ--KDHDKKDA-GT------------------   144
AtMYC3        ISSV------ENQNR-----------QSSCLVEKDLTFQGGLL-----------------------KSNETLSFCG--------------NESSKKRT-SVSKGSNND-EGMLSF--STV   368
AtMYC4        SVEN------PNPKV-----------LKSCEM---VNFKNGIE-----------------------NG----------------------QEEDSSNK-KRSPVSNNE-EGMLSF--TSV   369
AtMYC2        ITENPNLDPTPSPVH-----SQTQNPKFNNTFSRELNFSTS----------SS-TLVKPRSGEILNFGDEGKRSSGNPDPSSYSGQTQFEN---KRKR-SM------------------   399
VviMYC2       LTENPSIMHNPQQQQ---------QIHTQGFFTRELNFSEFGFDGNNGRNGNL-HSLKPESGEILNFGDSKRSSC-SANGNMFSGHSQVVAEENKKRR-SPTSRGSAE-EGMLSF--TSG   422

AtbHLH03      ----------QAIGG-----SNQV---------------YGYEQGKDETLYLTDEQKPRKRGRKPANGREEALNHVEAERQRREKLNQRFYALRAVVPNISKMDKASLLADAITYITDMQ   366
AtbHLH013     RLLPPAQMQIDFSAASSRASENNSD------------GEGGGEWADAVGADESGNNRPRKRGRRPANGRAEALNHVEAERQRREKLNQRFYALRSVVPNISKMDKASLLGDAVSYINELH   479
AtbHLH017     --------QIEFAGSSVAASSNPSTNTQQEKSESCTEKRPVSLLAGAGIVSVVDEKRPRKRGRKPANGREEPLNHVEAERQRREKLNQRFYALRSVVPNISKMDKASLLGDAISYIKELQ   441
AtbHLH014     ------------------------------------ETTRVAA------------------ATKEKHHPAVLSHVEAEKQRREKLNHRFYALRAIVPKVSRMDKASLLSDAVSYIESLK   295
VvibHLH009    --------TVG----RSSSDSGHSD-----------SDEPFASA-------LTENIRPKKRGRKPATGREMPLNHVEAERQRREKLNHRFYALRAVVPNVSRMDKASLLADAVSYIHELK   234
AtMYC3        VR-------------SAANDSDHSD-----------LEASVVKE---AIVVEPPEKKPRKRGRKPANGREEPLNHVEAERQRREKLNQRFYSLRAVVPNVSKMDKASLLGDAISYINELK   461
AtMYC4        LP-------------CD---SNHSD-----------LEASVAKEAESNRVVVEPEKKPRKRGRKPANGREEPLNHVEAERQRREKLNQRFYSLRAVVPNVSKMDKASLLGDAISYISELK   462
AtMYC2        VLNEDK--VLSFG-DKTAGESDHSD-----------LEASVVKE-------VAVEKRPKKRGRKPANGREEPLNHVEAERQRREKLNQRFYALRAVVPNVSKMDKASLLGDAIAYINELK   498
VviMYC2       VILPSS--CVVKS-SGGGGDSDHSD-----------LEASVVREADSSRV-VEPEKRPRKRGRKPANGREEPLNHVEAERQRREKLNQRFYALRAVVPNVSKMDKASLLGDAISYINELR   527
                                                                                    . :   *.*****:*******:***:**::**::*:*******.**::**  .::

AtbHLH03      KKIRVYETEKQIMKRRESN--------------------------------QITPAEVDYQQRHDDAVVRLSCPLETHPVSKVIQTLRENEVMPHDSNVAITEEGVVHTFTLRPQGGCTA   454
AtbHLH013     AKLKVMEAERERLGYS------------------------------SNPPISLDSDINVQTSGEDVTVRINCPLESHPASRIFHAFEESKVEVINSNLEVSQDTVLHTFVVKSE-----   563
AtbHLH017     EKVKIMEDERVGTDKSLSES---------------------------NTITVEESPEVDIQAMNEEVVVRVISPLDSHPASRIIQAMRNSNVSLMEAKLSLAEDTMFHTFVIKSNNGSD-   533
AtbHLH014     SKIDDLETEIKKMKMTETDKL---DNSSSNTSPS---SVEYQVNQKPSKSNRGSDLEVQVKIVGEEAIIRVQTENVNHPTSALMSALMEMDCRVQHANASRLSQVMVQDVVVLVPEGLR-   408
VvibHLH009    TKIDDLETKLREEVRKPKACLAEMYDNQSTT------TTSIVDHGRSSSSYGAIRMEVDVKIIGSEAMIRVQCPDLNYPSAILMDALRDLDLRVLHASVSSVKELMLQDVVVRIPEGLT-   347
AtMYC3        SKLQQAESDKEEIQKKLDGMSKEGNNGKGCGSRA-----KERKSSNQDSTASSIEMEIDVKIIGWDVMIRVQCGKKDHPGARFMEALKELDLEVNHASLSVVNDLMIQQATVKMGSQFF-   575
AtMYC4        SKLQKAESDKEELQKQIDVMNKEAGNAKSS---V-----KDRKCLNQ-ESSVLIEMEVDVKIIGWDAMIRIQCSKRNHPGAKFMEALKELDLEVNHASLSVVNDLMIQQATVKMGNQFF-   572
AtMYC2        SKVVKTESEKLQIKNQLEEVKLELAGRKASASGGDMS--------SSCSSIKPVGMEIEVKIIGWDAMIRVESSKRNHPAARLMSALMDLELEVNHASMSVVNDLMIQQATVKMGFRIY-   609
VviMYC2       TKLQSAESDKEDLQKEVNSMKKELASKDSQYSGSSRPPPDQDLKMSNHHGSKLVEMDIDVKIIGWDAMIRIQCSKKNHPAAKLMGALKELDLDVNHASVSVVNDLMIQQATVKMGSRFY-   646
                *:    * .                                        ::: :    :. :*:      :* : .: :: : .    .:.    .: :.:  .:

AtbHLH03      EQLKDKLLASLSQ--------------------   467
AtbHLH013     ELTKEKLISALSREQTNSVQSRTSSGR------   590
AtbHLH017     PLTKEKLIAAFYPETSSTQPPLPSSSSQVSGDI   566
AtbHLH014     --SEDRLRTTLVRTLSL----------------   423
VvibHLH009    --SEESMRTAILKRMQ-----------------   361
AtMYC3        --NHDQLKVALMTKVGENY--------------   592
AtMYC4        --TQDQLKVALTEKVGECP--------------   589
AtMYC2        --TQEQLRASLISKIG-----------------   623
VviMYC2       --TQDQLRLALSSKFADSR--------------   663
```

Supplemental Figure 3. Alignment of amino acid sequences of IIId- and IIIe-bHLHs from *Arabidopsis thaliana* (At) and *Vitis vinifera* (Vvi) was performed using the Clustal Omega program (https://www.ebi.ac.uk/jdispatcher/msa/clustalo). Identical and similar amino acids are marked with asterisks and dots, respectively. The red box highlights the amino acid sequences corresponding to the activation domain in MYC2, with identical amino acids in this region shaded gray. The blue box indicates the bHLH domain.

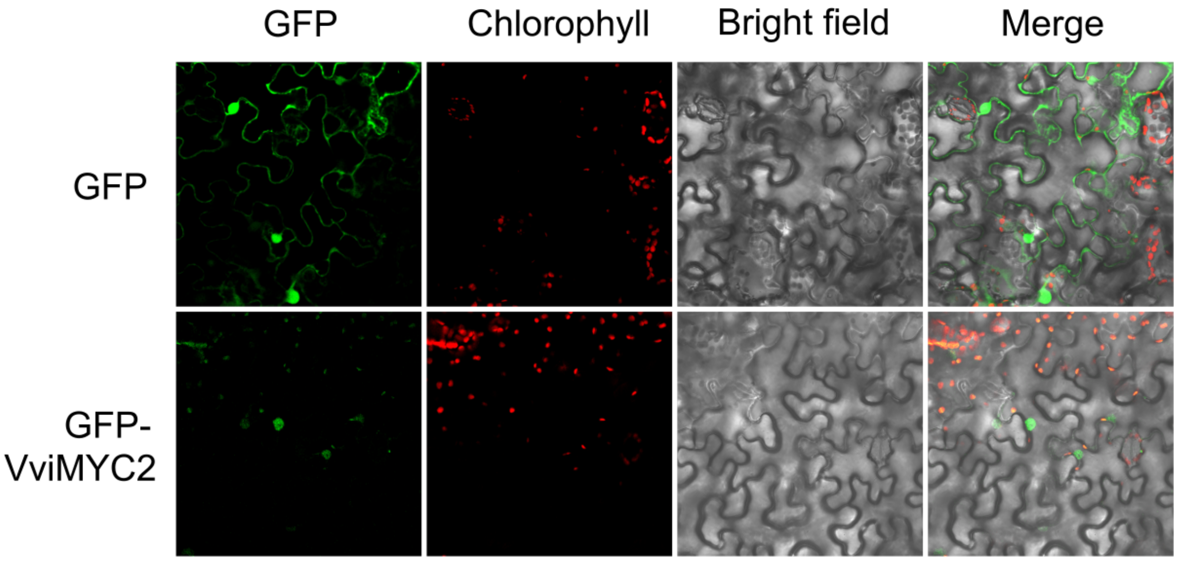


Supplemental Figure 4. The VviMYC2 transcription factor is localized in the nucleus. Its localization was demonstrated by the transient expression of a 35S:GFP-VviMYC2 construct in *Nicotiana benthamiana* leaves. Green fluorescence from GFP was observed two days after agroinfiltration, under 40× magnification (excitation at 488 nm, emission channel at 490–544 nm).

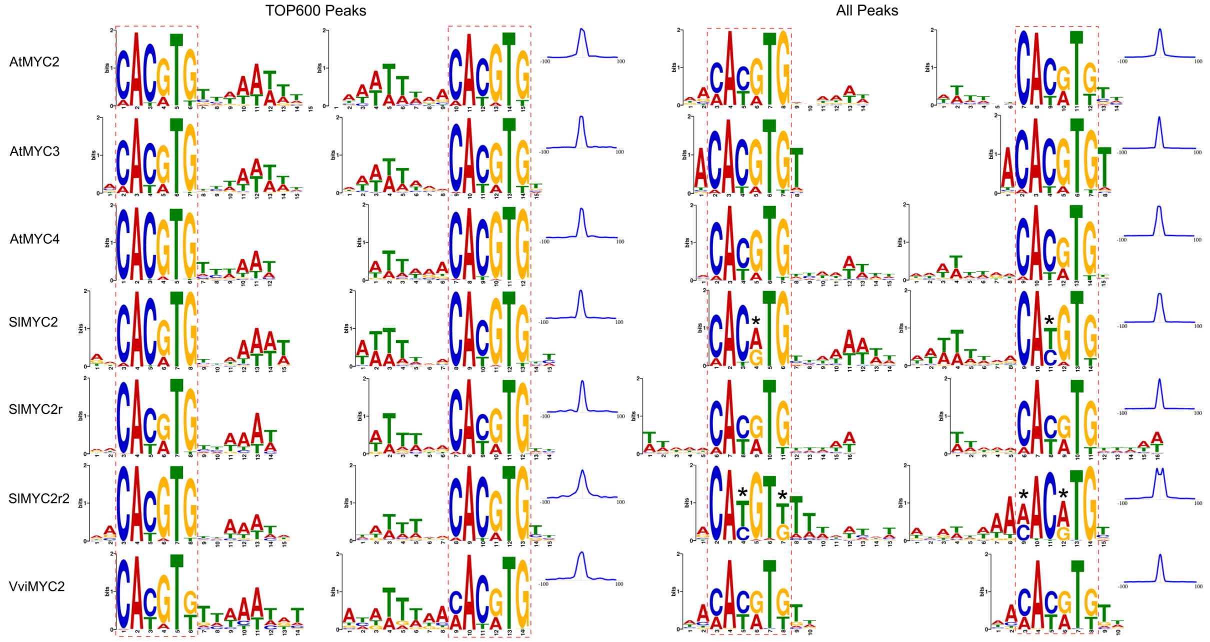


Supplemental Figure 5. Consensus DNA-binding motifs identified from DAP-seq data of IIIe-bHLH members in *Arabidopsis thaliana* (At), *Solanum lycopersicum* (Sl), and *Vitis vinifera* (Vvi) are shown. Binding motif hits were analyzed from the 600 most significant and all detected peaks of MYC2 DAP-seq, displayed for forward (left) and reverse complement (right) strands. The blue curve in the upper-right corner illustrates the binding motif profile within a 200 bp window frame of these peaks. Dashed pink boxes highlight the core sequences of consensus binding motifs, while asterisks indicate nucleotide differences in the core sequences compared to other motifs.

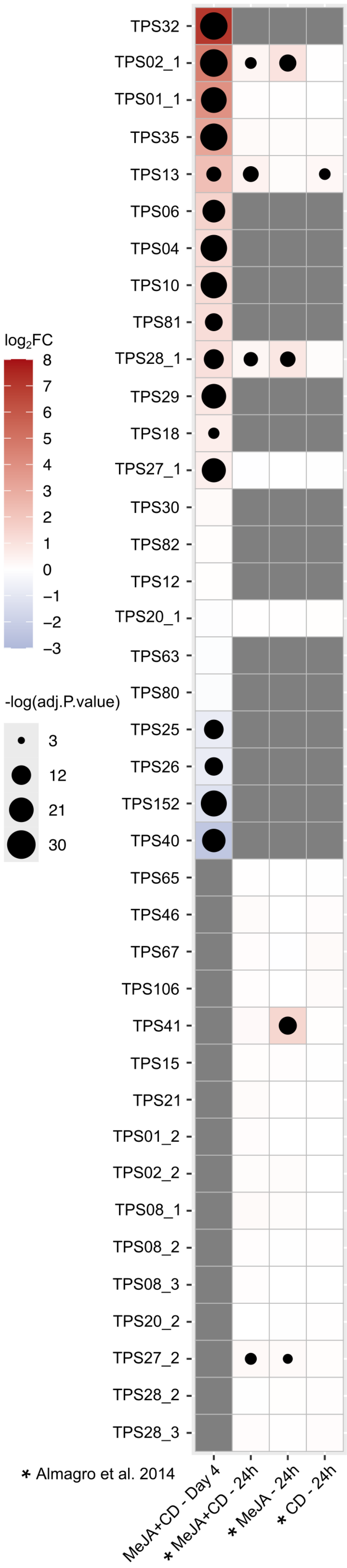
log2FC
8
7
6
5
4
3
2
1
0
−1
−2
−3
-log(adj.P.value)
3
12
21
30
TPS32
TPS02_1
TPS01_1
TPS35
TPS13
TPS06
TPS04
TPS10
TPS81
TPS28_1
TPS29
TPS18
TPS27_1
TPS30
TPS82
TPS12
TPS20_1
TPS63
TPS80
TPS25
TPS26
TPS152
TPS40
TPS65
TPS46
TPS67
TPS106
TPS41
TPS15
TPS21
TPS01_2
TPS02_2
TPS08_1
TPS08_2
TPS08_3
TPS20_2
TPS27_2
TPS28_2
TPS28_3
* Almagro et al. 2014
MeJA+CD - Day 4
* MeJA+CD - 24h
* MeJA - 24h
* CD - 24h

Supplemental Figure 6 (previous page). Differentially expressed *TPS* genes in MeJA- and MeJA+CD-elicited cells are presented. Statistical significance is depicted on a -log(adj. P-value) scale, indicated by dot size, while a color scale ranging from light blue to red represents differential expression relative to controls (log2FC). *TPS* genes targeted by multiple DNA probes on the microarray are marked with underscores and numbers.

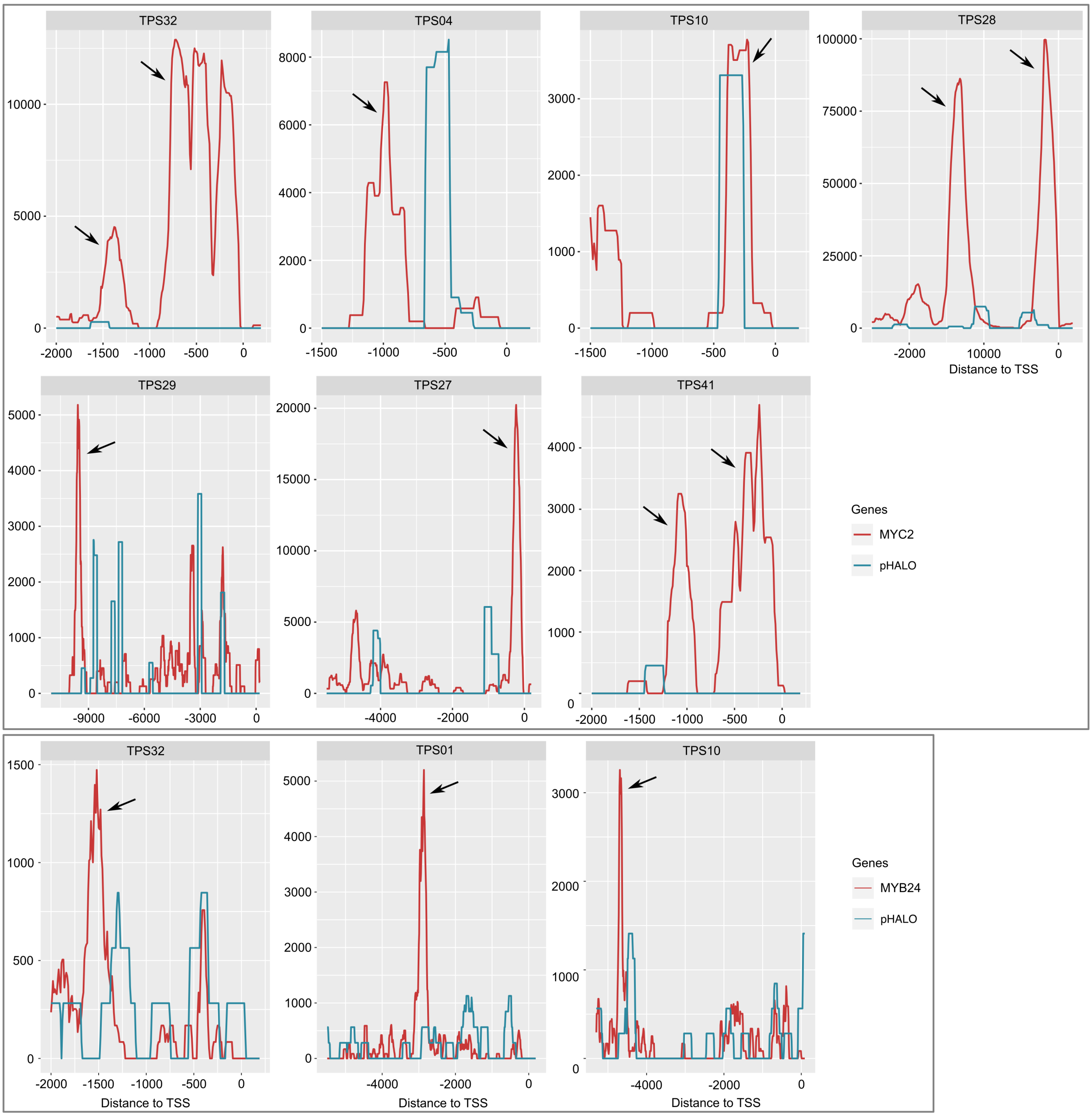


Supplemental Figure 7. MYC2 and MYB24 bind to the promoters of *TPS* genes. DAP-seq binding signals were compared to the empty vector control (pIX-HALO), with arrows indicating the detected binding peaks.

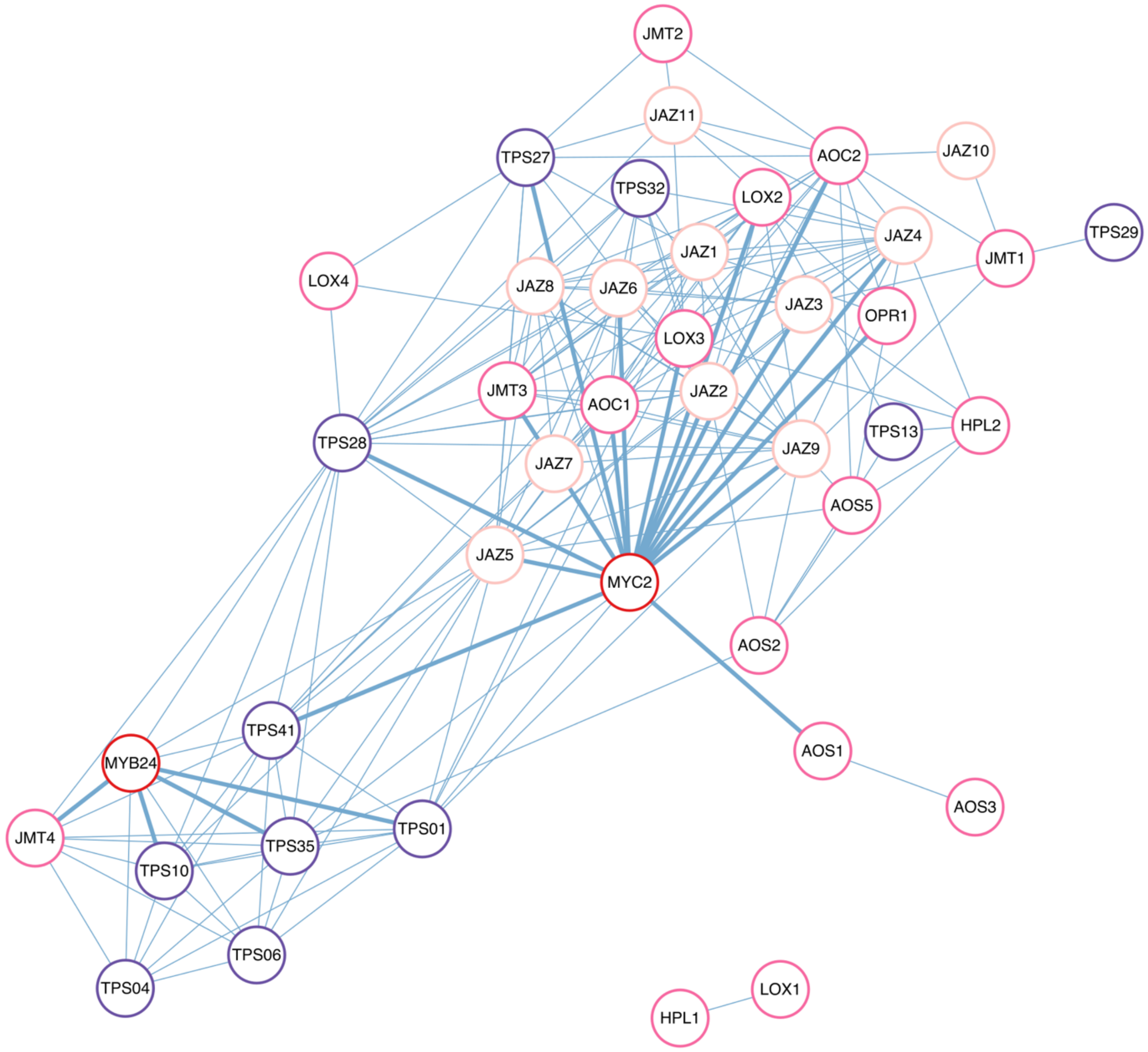


Supplemental Figure 8. Tissue-specific co-expression relationships between *MYC2*, *MYB24*, *TPSs* and jasmonate-related genes derived from fruit and flower aggregated co-expression networks (GCNs). Node distances were calculated using the d3Network package based on attraction metrics, which incorporate edge number and weight (default node repulsion mode). Co-expression metrics were queried and downloaded from the AggGCN app in the PlantaeViz platform (http://plantaeviz.tomsbiolab.com/vitviz).
